# Germ plasm anchors at tight junctions in the early zebrafish embryo

**DOI:** 10.1101/2021.03.24.436784

**Authors:** Nadia Rostam, Alexander Goloborodko, Stephan Riemer, Andres Hertel, Sabine Klein, Dietmar Riedel, Gerd Vorbrüggen, Roland Dosch

**Author notes:** Corresponding author: Correspondence to Nadia Rostam.

## Abstract

The zebrafish germline is specified during early embryogenesis by inherited maternal RNAs and proteins collectively called germ plasm. Only the cells containing germ plasm will become part of the germline, whereas other cells will commit to somatic cell fates. Therefore, proper localization of germ plasm is key for germ cell specification and its removal is critical for the development of soma. The molecular mechanism underlying this process in vertebrates is largely unknown. Here we show that germ plasm localization in zebrafish is similar to *Xenopus* and amniotes but distinct from *Drosophila*. We identified non muscle myosin II (NMII) and tight junction (TJ) components as interaction candidates of Bucky ball (Buc), which is the germ plasm organizer in zebrafish. Remarkably, we also found that TJ protein ZO1 colocalizes with germ plasm and electron microscopy (EM) of zebrafish embryos uncovered TJ like structures at early cleavage furrows. In addition, injection of the TJ-receptor Claudin-d (Cldn-d) produced extra germ plasm aggregates. Our findings discover for the first time a role of TJs in germ plasm localization.

## Introduction

Germ plasm consists of a maternally inherited ribonucleo-protein (RNP) condensate, which controls in many animals the formation of germline (Strome & Updike, 2015; Tristan Aguero, Susannah Kassmer, Ramiro Alberio, Andrew Johnson, 2017). Germ plasm thereby acts as a classical cytoplasmic determinant during embryonic development with the following activities: (I) In the zygote, germ plasm is asymmetrically localized, which after the cleavage period, leads to the formation of a subpopulation of embryonic cells containing germ plasm.. (II) These cells will be reprogrammed to differentiate into primordial germ cells (PGCs), while other cells without germ plasm adopt a somatic fate *e.g.* neuron, muscle etc. The correct positioning of germ plasm is therefore crucial for germline development, and its removal for the formation of somatic tissues. The molecular activities in germ plasm specifying PGCs seem to be largely conserved during evolution, because many components like Vasa, Nanos and Piwi are present throughout most animal genomes (Ewen-Campen et al., 2010; Juliano et al., 2010). By contrast, it is currently unknown whether the molecular mechanisms controlling localization of germ plasm is also conserved during evolution.

The positioning of germ plasm during embryogenesis is best understood in invertebrates, because of their powerful molecular-genetic tools. In *C. elegans*, the entry of sperm determines embryonic polarity (Otto & Goldstein, 1992; Strome & Wood, 1983), which eventually leads to asymmetric localization of germ plasm and germline specification (Seydoux, 2018; Strome & Updike, 2015). In the fly *Drosophila*, local translation of the germ plasm organizer Oskar (Osk) recruits germ plasm components to the posterior pole (Anne Ephrussi, 1992; Kim-Ha et al., 1993; Trcek & Lehmann, 2019). Among vertebrates using germ plasm for germline specification, some key discoveries of its localization were made in the frog *Xenopus laevis* (Tristan Aguero, Susannah Kassmer, Ramiro Alberio, Andrew Johnson, 2017).

For instance, a connection between the prominent Balbiani body (BB), also called mitochondrial cloud, and the germ plasm was first noticed in *Xenopus* (Heasman et al., 1984). Then, germ plasm gets anchored at the vegetal pole during oogenesis and after fertilization is passively inherited during the cleavage period at the forming furrows of the most vegetal blastomeres (Ressom & Dixon, 1988; Tristan Aguero, Susannah Kassmer, Ramiro Alberio, Andrew Johnson, 2017). At the blastula stage, germ plasm positive cells internalize into the embryo and then start their migratory journey until they reach the gonads. However, the molecular structure tethering germ plasm to the vegetal pole during the cleavage period of *Xenopus* embryogenesis is not known.

In zebrafish egg, germ plasm is also localized to the vegetal pole like in *Xenopus*, but this similarity of its positioning changes at the end of oogenesis (Dosch, 2015; Moravec & Pelegri, 2020; Raz, 2003). After fertilization, germ plasm streams together with cytoplasm during ‘ooplasmic segregation’ into the forming blastodisc at the animal pole (Welch & Pelegri, 2014). Subsequently, germ plasm localizes to the first two cleavage furrows, anchoring in four points at the four-cell stage (Olsen et al., 1997; Raz, 2003; Yoon et al., 1997). Indeed, maternal mutants affecting the first embryonic cleavages also interfere with germ plasm recruitment (Nair et al., 2013; Yabe et al., 2007). The first described cytoskeletal structure tethering germ plasm in zebrafish was described as furrow-associated microtubule-array (FMA) (Jesuthasan, 1998; Pelegri et al., 1999). However, the FMA starts to disassemble after the third cleavage, leaving the molecular identity of the cellular structure anchoring germ plasm after the eight-cell stage unresolved.

Molecular and genetic screens identified the first proteins which are specifically localized to these four spots, *e.g.* zebrafish Piwi (Ziwi)(Houwing et al., 2007), phosphorylated non muscle myosin II (p-NMII)(Nair et al., 2013) and Bucky ball (Buc) (Bontems et al., 2009; Campbell et al., 2015; Riemer et al., 2015; Roovers et al., 2018). Buc appears to exert a central role during germline specification, because it acts as a germ plasm organizer by recruiting other germ plasm components and thereby triggers germline specification (Bontems et al., 2009; Heim et al., 2014; Krishnakumar et al., 2018; Marlow & Mullins, 2008). Buc interacts through Kinesin Kif5Ba with microtubules, which is essential for Buc transport (Campbell et al., 2015). However, it is not clear, which cellular structure anchors Buc after its transport to the four spots in the early embryo.

We previously published that Buc and Osk have equivalent activities during germline specification (Krishnakumar et al., 2018) in zebrafish. Here, we show that the germ plasm nucleators Buc and its *Xenopus* homolog Velo1 use conserved mechanisms for their localization, whereas *Drosophila* Osk localizes by a distinct mode. We also show that Buc colocalizes with germ plasm in amniotes. We then mapped the localization motif in the Buc protein and used the isolated peptide to purify interactors from zebrafish embryos. Among numerous proteins, we identified subunits of the NMII complex, which is a known cytoskeletal component of adherens junctions, tight junctions and midbodies (Liu et al., 2012; Vicente-Manzanares et al., 2009). When we compared the localization of germ plasm with these cellular structures, we discovered that TJ protein ZO1 colocalizes with germ plasm. Electron microscopy (EM) of zebrafish embryos uncovered TJ like structures at those cleavage furrows that are in proximity to germ plasm at the 8-cell stage. Moreover, overexpressing the tight junction receptor Claudin-d (Cldn-d) led to the formation of ectopic germ plasm aggregates in zebrafish embryos. Taken together, our results indicate TJ as the cellular structure which recruits germ plasm at the onset of zebrafish embryogenesis.

## Results

### Zebrafish Buc and Xenopus Velo1 localize similarly in zebrafish embryos

The germ plasm organizers zebrafish Buc, *Xenopus* Velo1 and *Drosophila* short Oskar (sOsk) share the remarkable ability to specify germ cells in zebrafish (Krishnakumar et al., 2018). Consistent with this function as a germ plasm organizer, Buc localizes to the germ plasm (Bontems et al., 2009; Heim et al., 2014; Riemer et al., 2015) throughout early embryogenesis. To address whether this localization mechanism is also conserved between zebrafish, *Xenopus* and *Drosophila*, we injected mRNA encoding GFP-fusions of these germ plasm organizers into 1-cell zebrafish embryos (Fig. 1A). At 2.5-3 hours post fertilization (hpf), we compared the localization of the GFP-fusion proteins to the germ plasm using an antibody against the endogenous Buc proteins, which is tightly associated with the germ plasm (Bontems et al., 2009) and an antibody detecting β-Catenin to label membranes. Western blots with *in vitro* translated proteins confirmed that the Buc antibody did not cross-react with GFP, sOsk or Velo1 and thus specifically highlights endogenous germ plasm (Supplementary Fig. 1). Injections of mRNA encoding Buc-GFP colocalized with zebrafish germ plasm recapitulating the positioning of endogenous Buc (Fig. 1B, C, Supplementary Fig. 2A) (Bontems et al., 2009). Similarly, Velo1-GFP colocalized with the germ plasm (Fig. 1B, D, Supplementary Fig. 2B), suggesting that vertebrate Buc and Velo1 are targeted by a similar molecular machinery for germ plasm localization. By contrast, sOsk-GFP did not overlap with the germ plasm in injected zebrafish embryos (Fig. 1B, E, Supplementary Fig. 2C), but instead localized to the nuclei as previously shown in insect cells and *Drosophila* embryos (Jeske et al., 2017; Kistler et al., 2018). In contrast, injection of a GFP-control resulted in a ubiquitous subcellular localization including the nucleus (Fig. 1B, F, Supplementary Fig. 2D). These results suggest that zebrafish Buc and *Xenopus* Velo1 are localized by a conserved machinery, which does not recognize *Drosophila* Osk.

**Figure 1.**
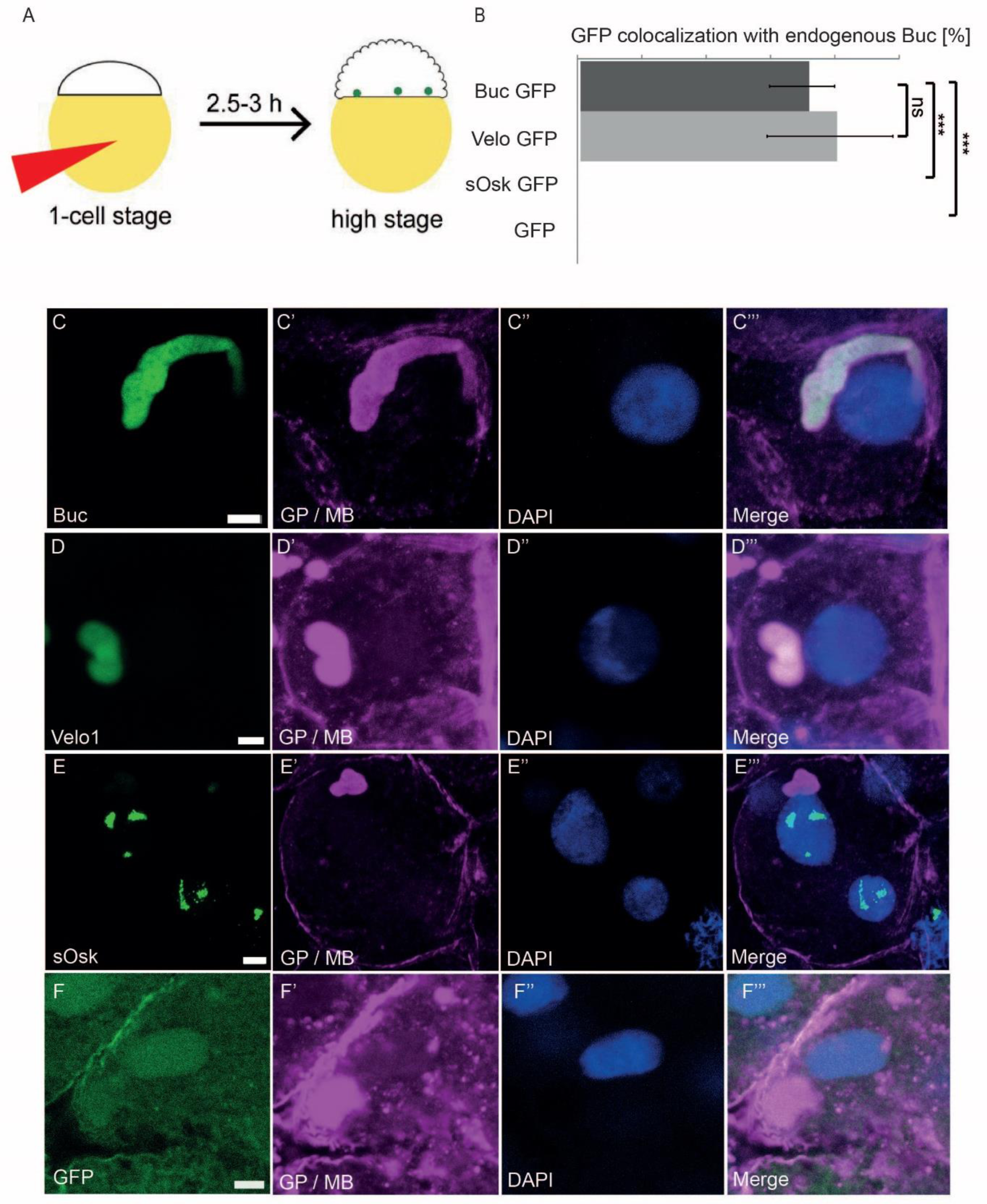
Buc and Velo1, but not *Drosophila* Oskar localize to zebrafish germ plasm. (A) Scheme of zebrafish colocalization assay. RNA encoding GFP fusions of germ plasm organizers Bucky ball (Buc), Velo1 and short Oskar (sOsk) was injected into 1-cell stage and scored at high stage for localization with endogenous Buc (green dots) by immunohistochemistry. (B) Quantification of colocalization assay. GFP fusions of Buc (71±10.1%) and Velo1 (79.7±19.5%; p=0.6), but not sOsk (0%; P=0.0005) or GFP alone (0%; P=0.0005) show colocalization with endogenous Buc. (C-F) Magnified germ plasm spot of embryo at high stage (full embryos are shown in the Supplementary Fig. 2). Colocalization of GFP (1st column, green) with endogenous Buc (germ plasm, GP; and -catenin to label membranes, MB; magenta) and nuclei (DAPI, blue) was determined by immunohistochemistry. n (Buc: 33, Xvelo:39, sOSK: 25, GFP: 32). Error bars represent standard deviation (SD). Scale bars: 5 µm.

### Buc does not localize in Drosophila embryos

The distribution of Osk protein in *Drosophila* is controlled by localization of *osk* mRNA to the posterior pole in the oocyte. This results in an exclusive localization of Osk protein to the posterior pole of the embryo, where it is eventually recruited to the centrosomes of forming pole cells (Lehmann, 2016; Lerit & Gavis, 2011). The recruitment of germ plasm and the formation of germ cells can be induced ectopically by targeting Osk translation to the anterior pole (Anne Ephrussi, 1992). We used this approach to address whether Buc is targeted by the machinery that localizes Osk in the *Drosophila* embryo. We fused the *buc* ORF to GFP and a *bicoid*-3’-UTR to direct its translation to the anterior pole of *Drosophila* embryos (Fig. 2A). As a positive control, we used *short osk* ORF fused to *bicoid*-3-UTR (*sosk*) (Tanaka & Nakamura, 2008). Indeed, immunolabelling of stage 4-5 fly embryos showed that Osk-GFP is anchored at the anterior cortex of the embryos and around the anterior nuclei of ectopically induced PGCs (Fig. 2B, D). By contrast, Buc was neither localized to the embryo cortex nor did it form perinuclear foci (Fig. 2E). Although the strength of the Buc-GFP signal increased during the onset of cellularization (stage 5), the protein was not anchored to the anterior cortex but distributed in a gradient originating at the anterior pole (Fig. 2C, E). Furthermore, although Buc-GFP was expressed in the anterior pole of the embryos, no cells were formed that showed morphological features of ectopic PGCs (Fig. 2C, E). Indeed, Vasa protein or *pgc* mRNA labeling in these transgenic embryos confirmed that sOsk specified ectopic PGCs anteriorly, whereas Buc-GFP did neither recruit Vasa protein or *pgc* mRNA to the anterior pole nor caused the formation of ectopic PGC (Supplementary Fig. 3). These results show that Buc is not recognized by the localization machinery in *Drosophila* that anchors sOsk to the cortex, suggesting that zebrafish and flies use different mechanisms for germ plasm localization.

**Figure 2.**
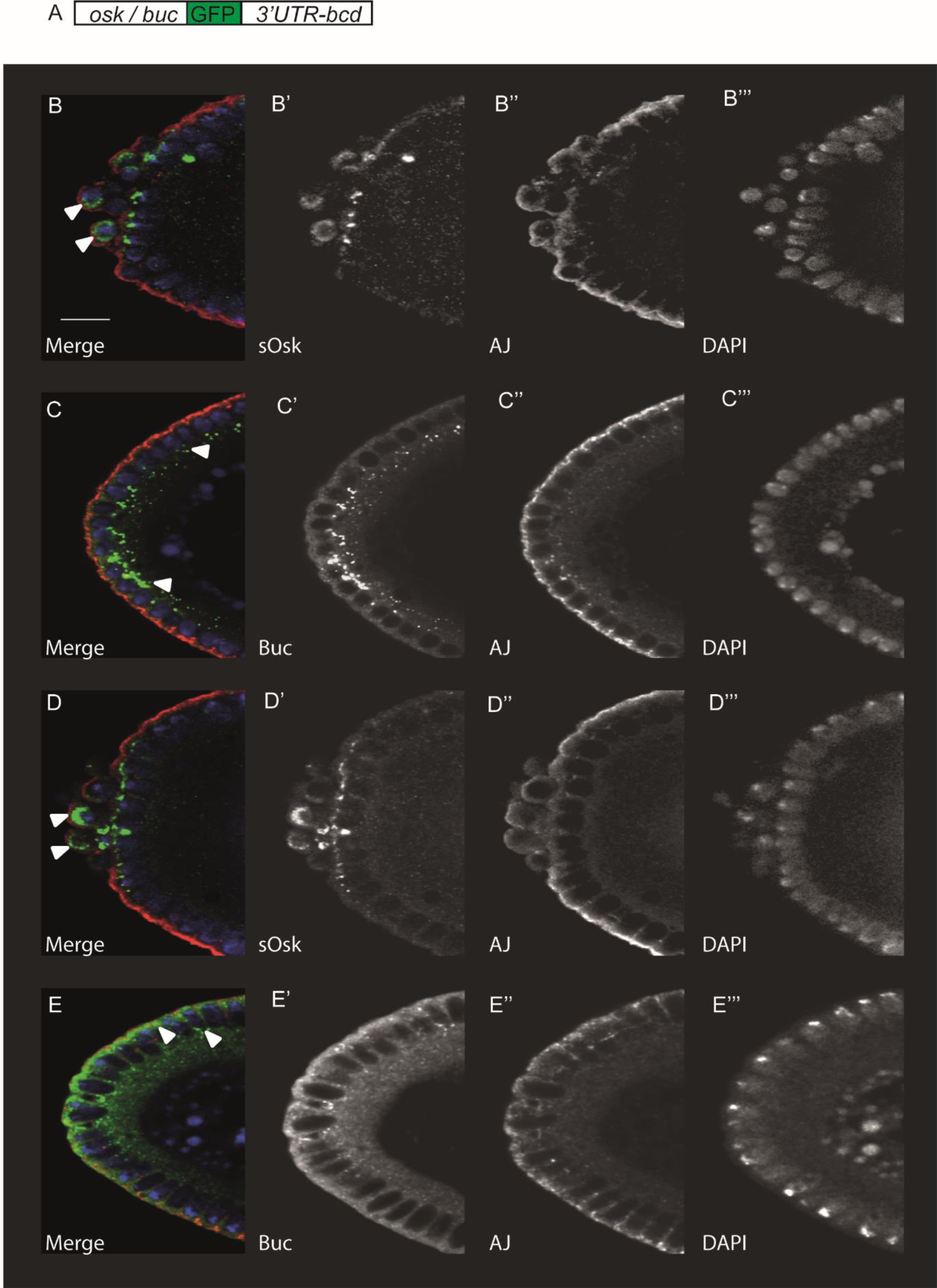
Transgenic Buc- and Oskar-GFP Drosophila embryos show different localization patterns. (A) Scheme of transgenes to study Bucky ball (Buc) and short Oskar (sOsk) localization. Transgenic flies were generated expressing Buc-GFP or sOsk ectopically at the anterior pole of the embryo by fusion of the constructs to the *bicoid* 3’UTR. (B-E) Localization of sOsk and Buc-GFP was investigated by immunohistochemistry: 1st column – merge, 2nd column –expressed protein, 3rd column – apical junctions (AJ), 4th column – DAPI. Anterior pole of immunostained embryos expressing the indicated transgenic constructs at stage 4 (B, C) and 5 (D, E). sOsk (B, B’, D, D’) localizes in condensed aggregates at the most distal part of the anterior pole (white arrowheads), whereas Buc-GFP (C, C’, E, E’) distributes in a gradient along the cortex of the anterior pole (white arrowheads). Scale bar: 10 µm.

### Buc localizes to the germ plasm in amniotes

Our results showed that zebrafish Buc and its functional *Xenopus* homolog Velo1 are localized to the germ plasm in zebrafish embryos, whereas *Drosophila* Osk did not show colocalization. To investigate if the localization machinery of germ plasm organizers might be further conserved within the amniotes, Buc localization in chicken embryos was analyzed. Chicken primordial germ cells are specified already in the laid egg blastoderm before they are relocated to the germinal crescent by the progressing primitive streak. Furthermore, the chicken Vasa homolog (Cvh) as an evolutionary conserved germ plasm marker, is localized in the germinal vesicle of chicken oocytes, the cleavage furrows of the dividing zygote and the large granular PGCs in the central epiblast (Tsunekawa et al., 2000).

To analyze the localization of Buc in the germline of chicken embryos, we used whole-mount immunohistochemistry. Double labelling with Buc and Cvh antibodies shows colocalization of Buc with the germ plasm of cleavage stage I and stage II chicken embryos (Fig. 3A, B).

**Figure 3.**
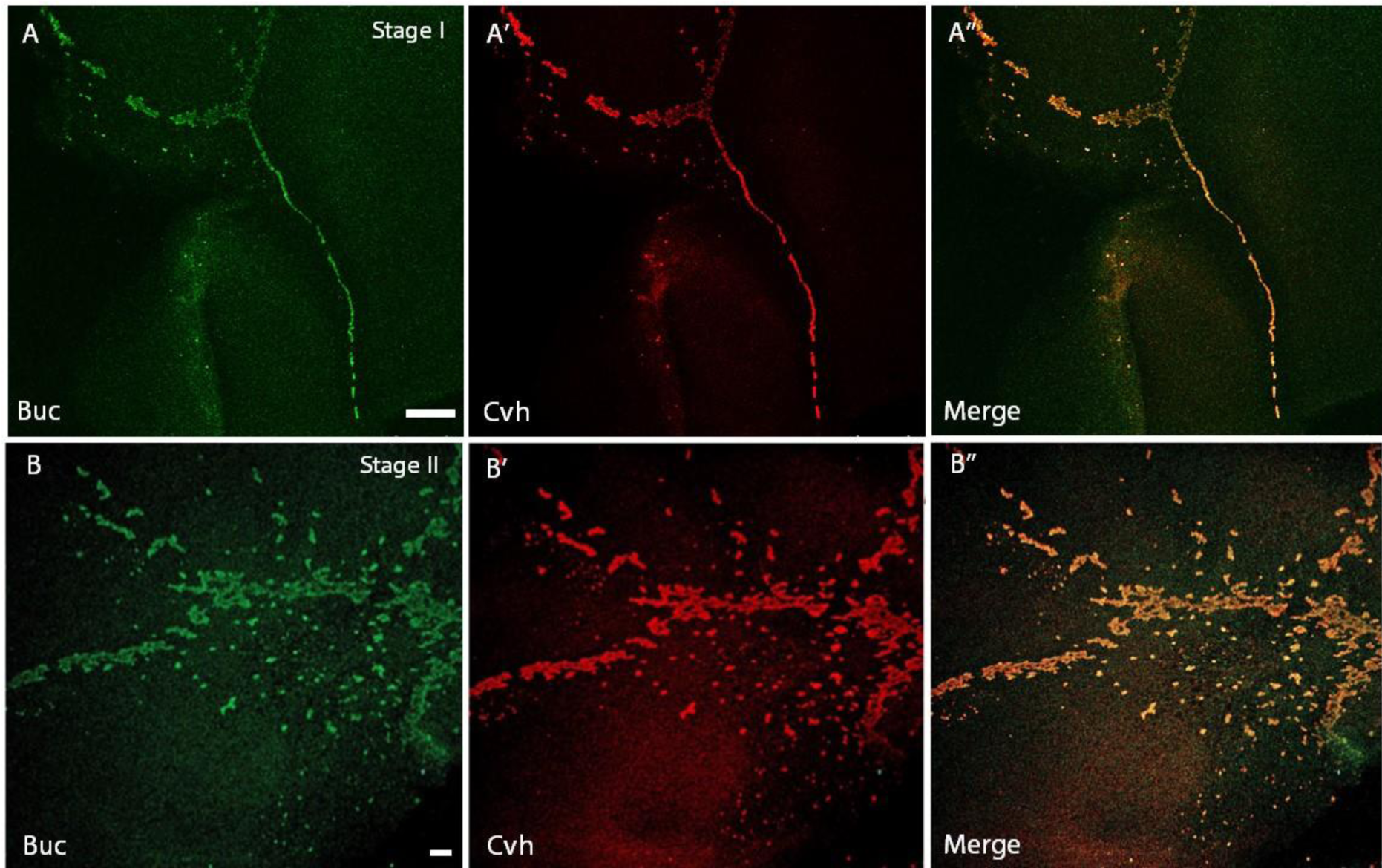
Buc localizes to the germ plasm in amniotes. Colocalization of Bucky ball (Buc) with the germ plasm marker Cvh (Vasa homolog) was determined by immunostaining in early chicken embryos. (A) Colocalization at stage I of embryonic development (3 hpf). (B) Colocalization at stage II of embryonic development (4 hpf). 1^st^ column - Buc (green), 2^nd^ column - Cvh (red), 3^rd^ column - merge (yellow). Note the strong colocalization of both proteins at the cleavage furrows and other granule scattering around the central furrows, as shown in the overlay at both stages (A’’ and B’’). Scale bars (A): 100 µm, (B): 20 µm.

To confirm the positioning of Buc at cleavage furrows, we co-labeled embryos with the membrane marker pan-Cadherin at stage II. Buc co-localizes with pan-Cadherin suggesting that germ plasm colocalizes with the membrane in the chicken embryo (Supplementary Fig. 4). Nonetheless, the more widespread distribution of pan-Cadherin implies that Buc localizes to a restricted region of the plasma membrane. Taken together, these results indicate that germ plasm localization to the membrane is evolutionary conserved between zebrafish and chicken embryos.

### The Buc localization signal is part of the conserved N-terminal BUVE motif

We have previously shown that the 3’-UTR of the *buc* mRNA is not required for its localization (Bontems et al., 2009). Therefore, we analyzed its amino acid sequence to identify the protein domain of Buc that is essential for the localization to the four germ plasm spots at 3 hpf. We generated systematic deletions of Buc fused to GFP (schematically shown in Fig. 4A), injected the mRNA into zebrafish 1-cell stage zygotes and scored the number of embryos with GFP foci at 3 hpf as depicted in Fig. 1A.

**Figure 4.**
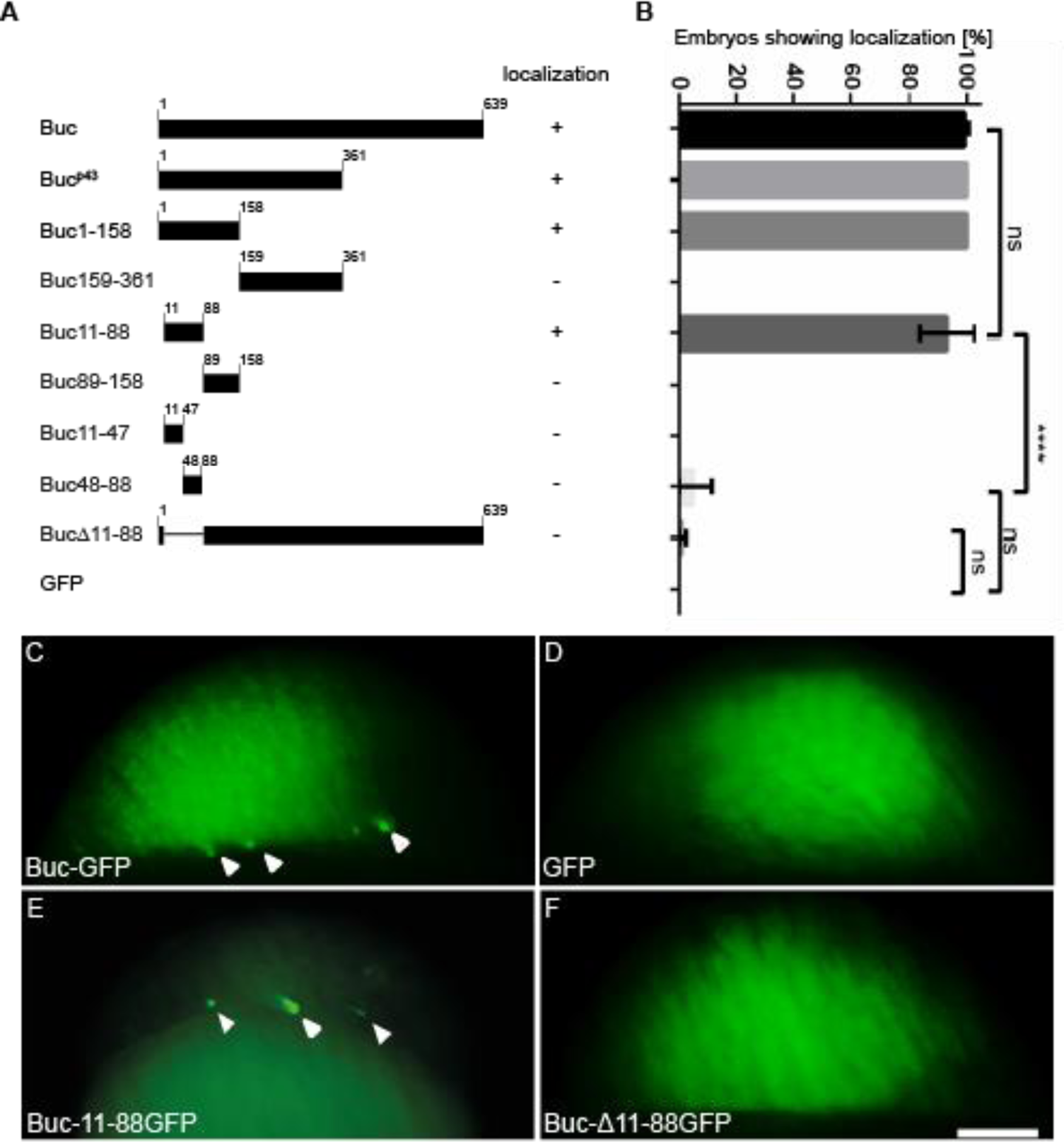
Buc11-88 is necessary and sufficient for Buc localization in zebrafish embryos. In this localization assay RNA was injected into 1-cell stage embryos and scored at high stage for localization of fluorescence in living embryos as shown in Fig. 1A. (A) Schematic representation of Buc protein deletions and summary of their localization (+/-). Numbers indicate amino acids. (B) Quantification of the localization assay. Buc11-88 localized (90.9±10.1%) similarly to WT Buc (99.1±1.3%) (P=0.8). BucΔ11-88 did not localize (0.9±1.6%) compared to WT Buc (P=0.009) and Buc11-88 (P=0.01. (C-F) Blastomeres of living high stage embryos oriented as shown in Figure 1A expressing the indicated constructs. Note protein localization of Buc-GFP (D; arrowheads) or Buc11-88 (F; arrowheads), whereas a GFP-control or BucΔ11-88 show ubiquitous fluorescence (E, G). n (Buc-GFP: 98, GFP: 94, Buc-11-88GFP: 181, Buc-Δ11-88: 230). Error bars represent SD. Scale bar: 100 µm.

An N-terminal fragment (aa 1-361, Fig. 4A) which corresponds to the previously identified *buc^p43^* mutant allele localizes correctly and with the same penetrance as full-length Buc (Fig. 4A, B, C, Supplementary Fig. 5A). Next, we split this fragment into two halves (aa1-158 and 159-361) and analyzed their localization. Buc1-158 localized, whereas Buc159-361 showed ubiquitous fluorescence, similar to control embryos injected with GFP mRNA (Fig. 4A, B, Supplementary Fig. 5B, C). We then split Buc1-158 into two fragments and in addition removed the first ten amino acids (Buc11-88), which show a low conservation in teleost evolution (Škugor et al., 2016). Buc11-88 was sufficient to recapitulate germ plasm localization, whereas Buc89-158 showed no specific localization (Fig. 4A, B, E, Supplementary Fig. 5D). Further splitting of Buc11-88 disrupted the localization activity of both resulting fragments (Fig. 4A, B, Supplementary Fig. 5E, F), suggesting that aa 11-88 contains the residues sufficient to target the protein to the germ plasm spots. To confirm that Buc does not contain other motifves involved in localization, we generated a deletion of the isolated motif aa11-88 (BucΔ11-88) in full length Buc. This protein did not localize (Fig. 4A, B, F). We therefore concluded that aa11-88 is sufficient and necessary for the localization of Buc and named the protein-region BucLoc.

### Prion-like domains in the BucLoc motif are not required for Buc localization

Previous studies showed that the N-terminus of Buc including BucLoc is strongly conserved during vertebrate evolution (Boke et al., 2016; Bontems et al., 2009; Krishnakumar et al., 2018). This stretch of 100 amino acids, termed BUVE motif (for Buc-Velo), was recently shown to be responsible for Velo localization to the BB during *Xenopus* oogenesis (Boke et al., 2016). Furthermore, the localization of Velo1 to the BB is driven by aggregation of two prion-like domains (PLDs) within the BUVE motif (Boke et al., 2016). A Sequence alignment of Buc with Velo1 showed the conservation of the aromatic amino acids of the PLDs in Buc between aa24-30 and 64-71 (Fig. 5A, marked in red), suggesting that Buc might also have the ability to form amyloid-like assemblies. Therefore, we tested if the same aggregation mechanism might control the localization of Buc in the early zebrafish embryos.

**Figure 5.**
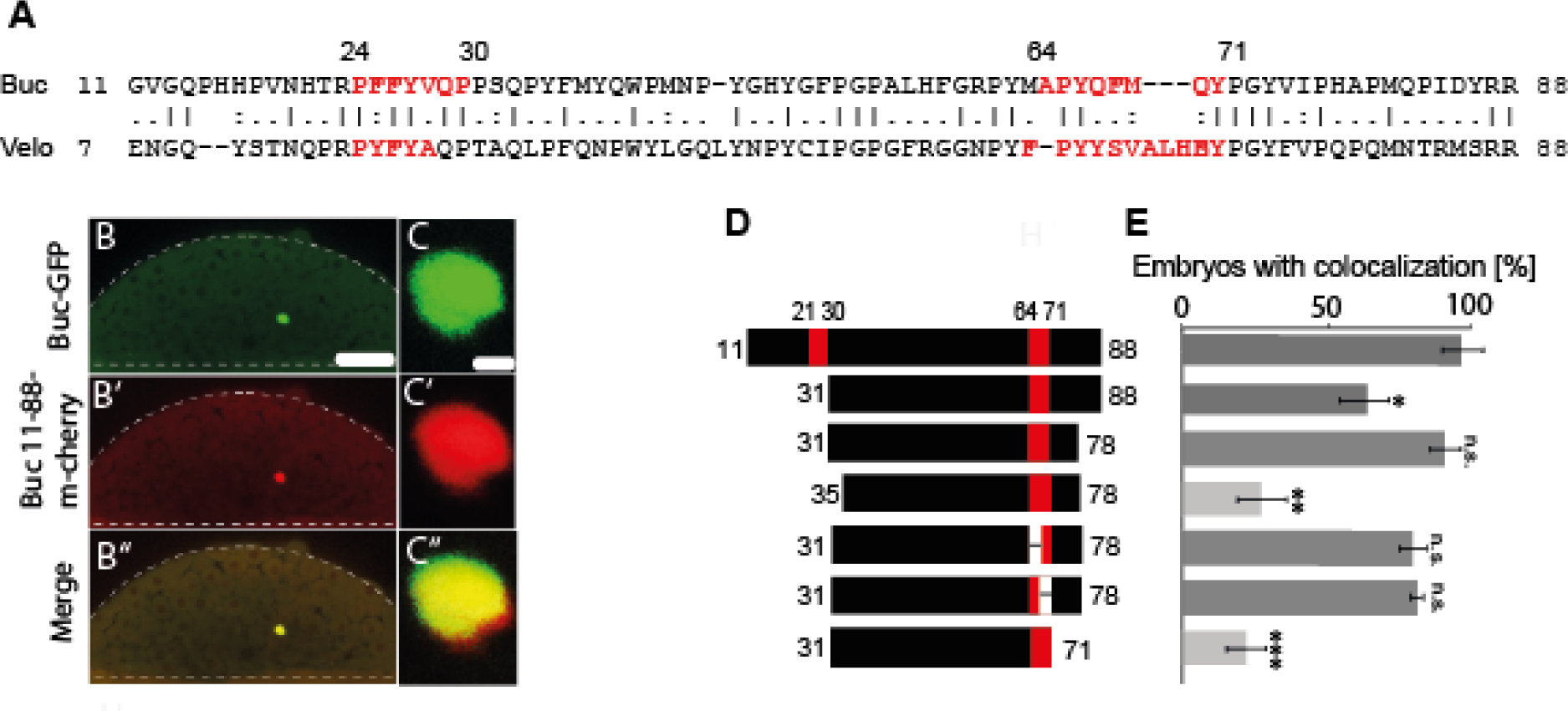
Aggregation and localization of BucLoc are separate activities. (A) Alignment of Buc11-88 with the N-terminus of *Xenopus* Velo1 (aa7-88). Red letters highlight the prion-like domains previously discovered in Velo1 (Boke, 2016) and their corresponding amino acids in Buc. (B-B’’, C-C’’) BucLoc (11-88)-m-cherry colocalizes in transgenic embryos with endogenous Buc-GFP to the germ plasm. (B-B’’) Living sphere stage transgenic Buc-GFP embryo injected at 1-cell stage with RNA encoding BucLoc-m-cherry, showing colocalization. Embryo is shown on the lateral view with the animal pole to the top, outlined by the white dashed line. (C-C’’) Magnification of the localized spot of germ plasm shown in (B-B’’). (D) Summary of BucLoc mapping showing that PLDs are not important for the localization of Buc. Prion-like domains are shown with red boxes. (E) Quantification of BucLoc mapping and 5aa deletions in (D). Buc31-88 (60.1± 7.9%) and Buc31-71 (21.1± 6.4) show significantly less localization compared to Buc11-88 (P= 0.01 and 0.0004). There was no significant difference between the localization of Buc11-88 and Buc 31-78 (P=0.41), excluding the role of the first PLD in localization. 5aa deletions of Buc31-78 showed that residues other than second PLD are important in the localization of Buc. Buc31-78 Δ 31-35 (30.0± 10) showed significantly less localization compared to Buc31-78 (P=0.009). Colocalization of constructs in (D) is shown in Supplementary figure 7. n (Buc11-88: 30, Buc31-88: 30, Buc31-78: 30, Buc35-78: 30, Buc31-78Δ62-66: 30, Buc31-78 Δ67-71: 30, Buc31-71). Error bars represent SD. Scale bars (B): 50 µm, (C): 2 µm.

To investigate the importance of these two potential PLD domains in Buc, the colocalization of deletion variants with the germ plasm was analyzed. Therefore, mRNA of deletion variants of the BucLoc domain fused to mCherry were injected into 1-cell embryos and the colocalization to germ plasm aggregates marked by Buc-GFP was examined at 3 hpf. As a positive control we used the entire BucLoc domain (aa11-88) that shows colocalization with the endogenous germ plasm (Fig. 5B, C, quantification in E). To narrow down the localization motif further, the N-terminal 20 amino acids were removed. Indeed, Buc 31-88 showed a slight reduction in germ plasm localization. Interestingly, when we deleted ten C-terminal amino acids (Buc31-78), localization was restored to nearly wild-type frequency (Fig. 5D, E, Supplementary Fig. 7B). By contrast, deleting four additional N-terminal amino acids (Buc35-78) almost completely abrogated localization. Nonetheless, the first PLD between aa25 and aa30 does not seem to be necessary for localization.

To examine the role of the second PLD, we generated internal deletions in Buc31-78. When we removed the second PLD (Δ64-71), no fluorescence could be detected in the embryos (Supplementary Fig. 7C) suggesting that aa64-71 might affect protein stability or translation. Therefore, we analyzed two variants with five amino acid deletions within the second PLD domain. Strikingly, removing parts of the second PLD (BucΔ62-66 or BucΔ67-71) caused no clear reduction in germ plasm localization (Fig. 5D, E, Supplementary Fig. 7D, E). In contrast, when we kept the second PLD domain intact, but removed C-terminal sequences (Buc31-71), the localization efficiency dropped to 15% (Fig. 5D, E, Supplementary Fig. 7F, Supplementary Fig. 8). These results suggest that the two PLD motifs, which control Buc’s aggregation into the Balbiani body during oogenesis (Boke et al., 2016), are not involved in positioning Buc to the germ plasm in the zygote.

### Identification of the BucLoc interactome

As Buc forms clusters with the germ plasm in the proximity of the cleavage furrows, we aimed to identify the cellular structure that is essential for its anchorage. One candidate is the furrow-associated microtubule-array (FMA), which was shown to be involved in tethering germ plasm in zebrafish (Jesuthasan, 1998; Pelegri et al., 1999). As our results show that BucLoc domain is sufficient for the localization of Buc to the germ plasm foci, we used this protein motif as a bait to identify cellular binding partners directly by co-immunoprecipitation followed by mass spectrometry analysis. Embryos were injected at 1-cell stage with mRNA encoding BucLoc-GFP, lysed at the stage of the formation of germ plasm foci and immunoprecipitated using GFP-tag (Fig. 6A). Embryos injected with mRNA encoding GFP were used as a negative control, and transgenic embryos for full length Buc-GFP were used to control for mRNA overexpression (Fig. 6A, B).

**Figure 6.**
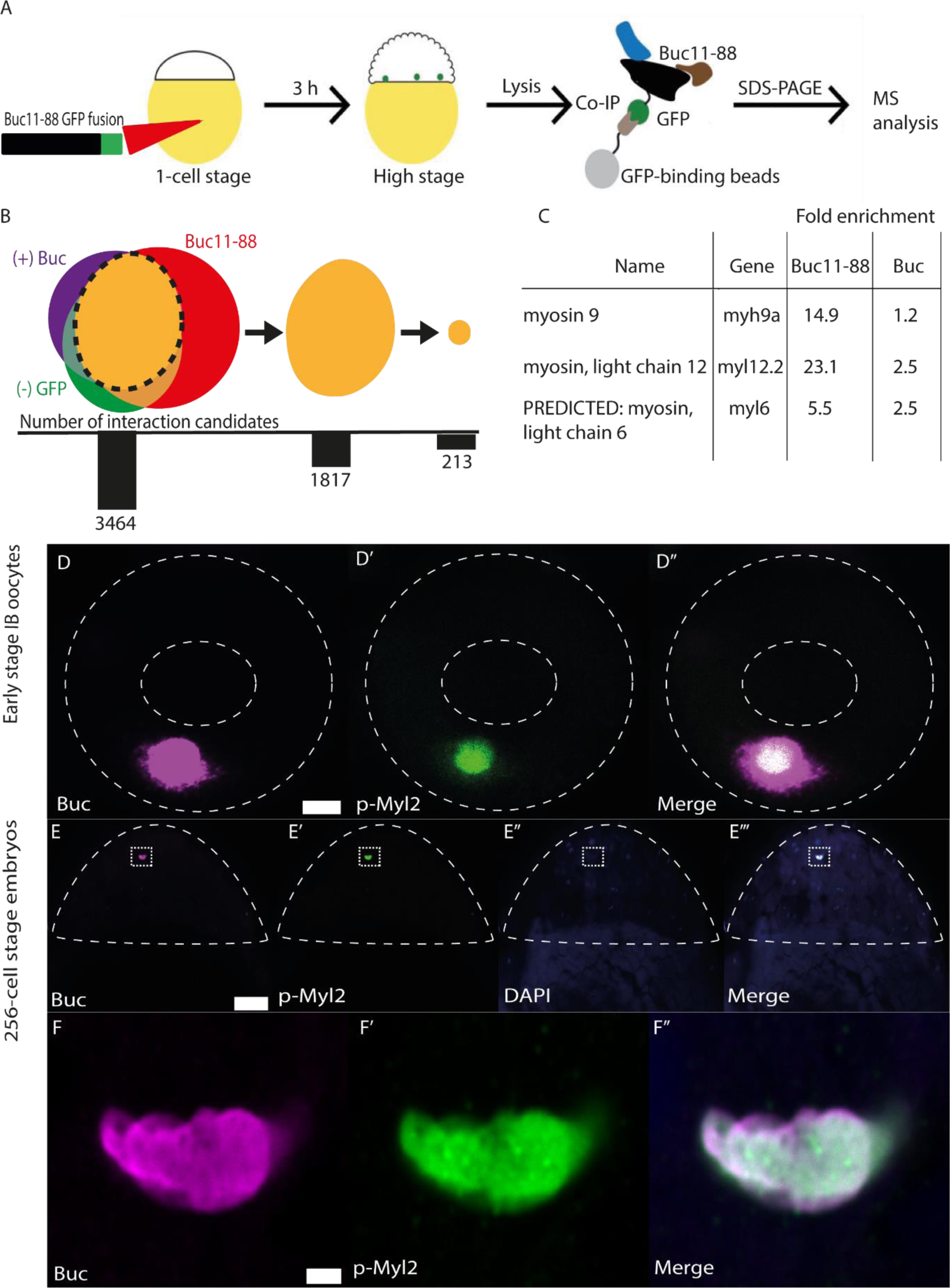
Non-muscle myosin II (NMII) colocalizes with Buc. Colocalization of Bucky ball (Buc) and phosphorylated myosin light chain 2 (p-Myl2) was determined by immunostaining. (A) Schematic representation of mass spectrometry of BucLoc. Wild type embryos (n: 500) were injected with RNA encoding for BucLoc-GFP and lysed at high stage. Embryos of the transgenic Buc-GFP line were used as positive control. GFP RNA injected embryos were used as negative control. Subsequent to lysis, an IP against the GFP-tag was carried out. Interacting proteins were identified by mass spectrometry. (B) 3464 proteins were identified in the mass spectrometry, of which 1817 candidates interacted with both Buc-GFP and BucLoc-GFP and 213 specifically with BucLoc-GFP (for selection criteria see Material and Methods). (C) Fold enrichment of myosin light chain in the mass spectrometry. (D-D’’) Colocalization at early stage (IB) oocyte stage. 1^st^ column -Buc (magenta), 2^nd^ column – p-Myl2 (green), 3^rd^ column - merge (white). (E-E’’) Show colocalization in embryo (256 cells). 1^st^ column - Buc (magenta), 2^nd^ column - Myl2 (green), 3rd column - DAPI (blue) and 4th column - merge (white). (F-F’’) Show magnification of germ plasm spot in E-E’’. Scalebars (D-D‘‘): 10 µm, (E-E‘‘‘): 50 µm, (F-F‘‘‘): 2 µm.

In this analysis, we found 1817 protein candidates that potentially interact with full length Buc and BucLoc but not with GFP. From those, 213 proteins were strongly enriched for BucLoc interaction (Supplementary Table 1 for the full list of candidates of the mass spectrometry) and represent therefore candidates for the subcellular network required for germ plasm localization. Among the candidates that were strongly enriched was Myosin Light Chain (Fig. 6C), which is a subunit of the Non-Muscle Myosin II (NMII) protein complex. Interestingly, phosphorylated NMII (p-NMII) colocalizes with germ plasm RNAs at the 2- and 4-cell stage in zebrafish embryos (Nair et al., 2013). To investigate if p-NMII also colocalizes with Buc and could therefore play a role in germ plasm localization we performed immunohistochemistry for Buc and p-NMII. Indeed, we found that Buc colocalizes with p-NMII in early stage IB oocytes (Fig. 6D) and during zebrafish embryogenesis (256 cell stage, Fig. 6E, F). These results confirm that our immunoprecipitation experiment has purified candidates involved in anchoring Buc in the forming germ plasm foci.

### Buc colocalizes with tight junction protein ZO1

NMII is known to associate with various cellular structures (Liu et al., 2012; Nair et al., 2013; Vicente-Manzanares et al., 2009). NMII is activated upon phosphorylation by various kinases including Rho-associated protein kinase (ROCK), which also regulate germ plasm compaction in the early zebra fish embryo (Amano et al., 1996; Miranda-Rodríguez et al., 2017). To identify the exact cytoskeletal component that anchors germ plasm, we looked at the structures that interact with NMII. p-NMII is part of adherens junctions (AJ) (Liu et al., 2012), the midbody (Wang et al., 2019) and tight junctions (TJ). To address which of these cellular structures could be involved in anchoring germ plasm, we compared the localization of Buc (Bontems et al., 2009) with marker proteins for tight junctions, adherens junctions and the midbody, respectively. We found that Buc abutted on adherens junctions (E-Cadherin, Fig. 8B, Supplementary Fig. 7B) and the midbody marker midbody (Kif23, Fig. 7C, Supplementary Fig. 9C). Fascinatingly, Buc colocalized with the TJ marker ZO1 (Fig. 7D, Supplementary Fig. 9A). These data show that Buc localizes to the ZO1-positive foci at the cleavage furrows suggesting that this cellular structure is responsible of the association of Buc to the FMA.

**Figure 7.**
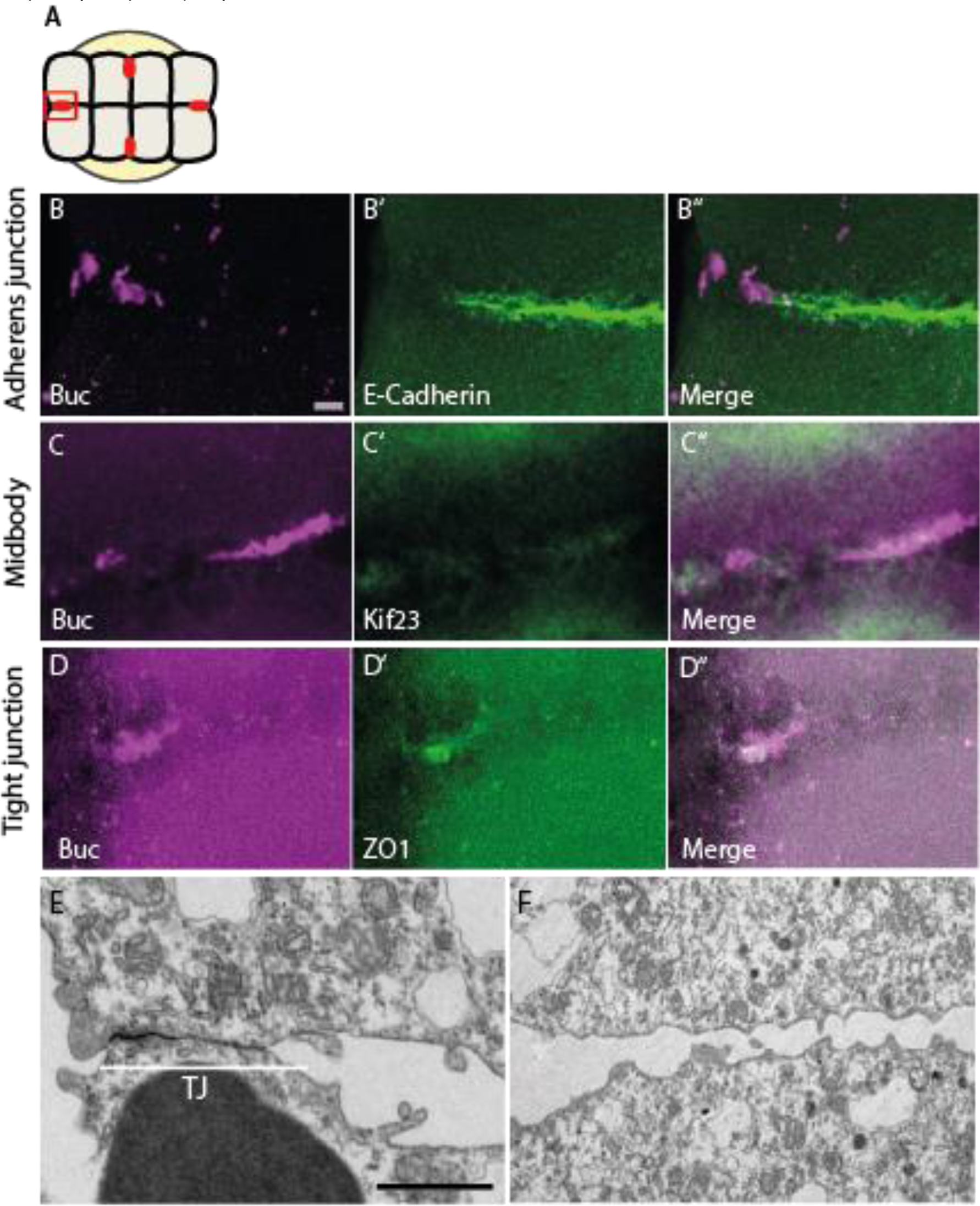
Buc colocalizes with TJ protein ZO1 and electron microscopy of early cleavage furrows shows TJ-like structures. Colocalization analysis of Buc with different cellular structure markers at 8-cell stage. (A) A representative cartoon showing an 8-cell stage embryo from the animal view. The red dots show where germ plasm is localized the cleavage furrows. The cleavage furrows which do not have red dotes do not contain germ plasm. The red box represent the cleavage furrows which are shown in the following pictures. (B, C, D) Magnification of one of the cleavage furrows containing germ plasm (full embryo staining is shown in Supplementary Fig. 7). 1^st^ column - Buc (magenta), 2^nd^ column – respective cellular structure (green), 3^rd^ column - merge. (B-B‘‘) Immunostaining for Buc and adherens junction marker E-cadherin; (C-C‘‘) Buc and midbody marker Kif23; (D-D‘‘) Buc and tight junction marker ZO1. (E) Electron microscopy of germ plasm containing cleavage furrow. Note: TJ-like structures are observed in this cleavage furrow as shown with the upper white line but no germ plasm granules are shown here. (F) Electron microscopy of a non-germ plasm containing cleavage furrow. Scale bars (B, C, D): 5 µm, (E, F): 1 µm.

**Figure 8.**
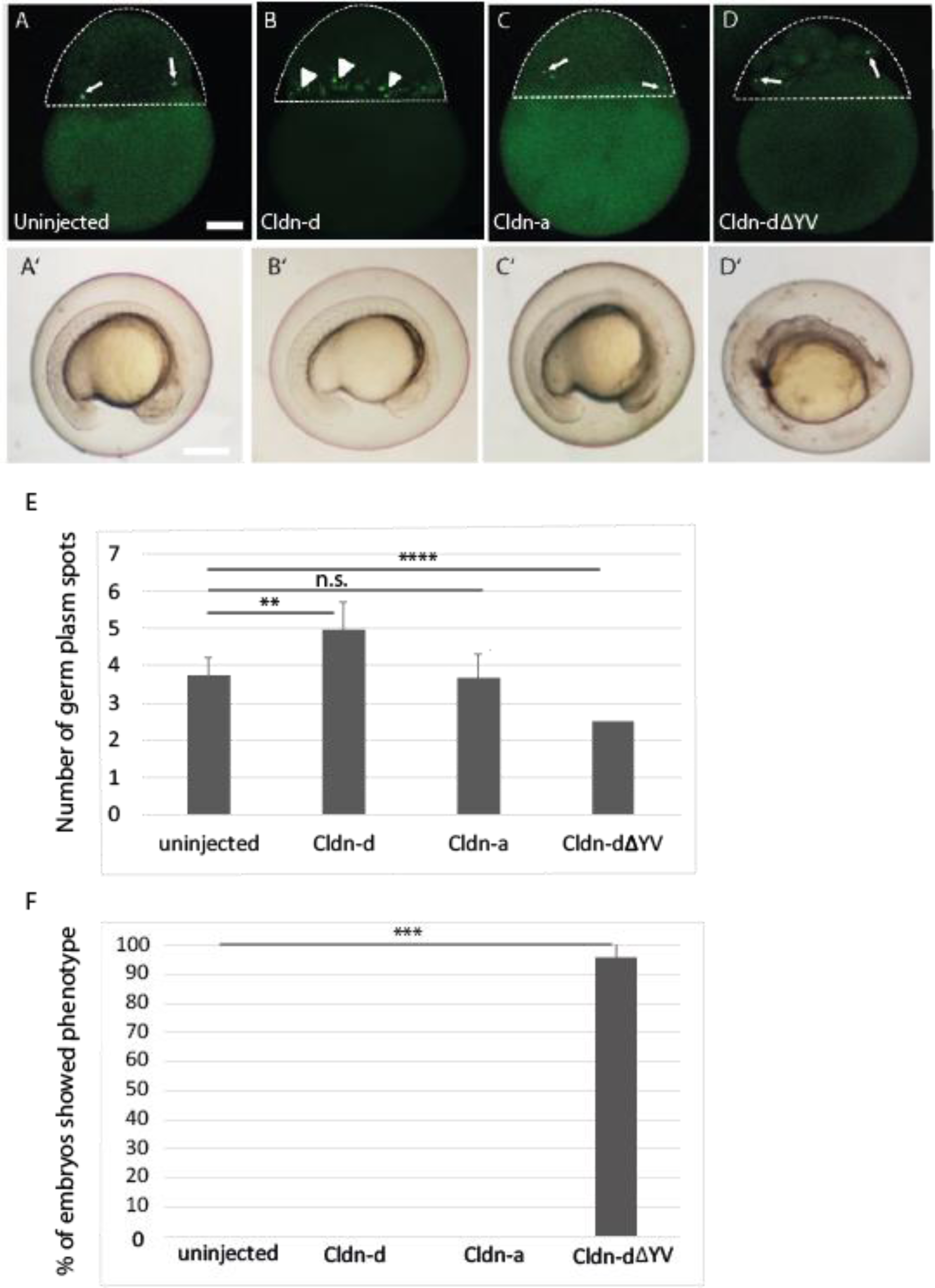
Cldn-d induces ectopic germ plasm foci. Over expression of Cldn-d in zebrafish embryos at 1-cell stage. A) An uninjected embryo from Buc-GFP transgenic line at 2hpf. Buc is localized in germ plasm spots. Arrows show germ plasm spots. B) Over expression of Cldn-d produces additional germ plasm spots (arrow heads). C) Embryo injected with c*ldn-a* show no effect in comparison to control (A). D) Injection of *cldn-dΔYV* produces a strong phenotype in zebrafish embryos. One embryo is shown here which has developmental defect and its blastomers are not properly attached to each other. Germ plasm spots are shown with arrows. A’-D’) Developed embryos from (A-D) at 1 dpf (day post fertilization). A’) An uninjected embryo, B’) A *cldn-d* injected embryo, C’) a c*ldn-a* injected embryo and D’) a *cldn-dΔYV* injected embryo showing developmental defects. E) Quantification of the average number of germ plasm spots in uninjected and injected embryos. c*ldn-a* injection showed no significant difference to uninjected control (P-value: 0.3), with average number of spots (3.66 ± 0.65) and (3.74 ± 0.48), respectively. *cldn-d* injection caused a significantly higher number of germ plasm spots (4.96 ± 0.76) compared to controls (P-value: 0.004) and *cldn-dΔYV* injection resulted in significantly lower number of germ plasm spots (2.49 ± 0.62) compared to controls (P-value: 0.00003) and c*ldn-a* injected embryos (P-value: 0.0049). Note germ plasm spots in A, B, and D (arrows) and ectopic germ plasm spots in C (arrow heads). F) Quantification of total number of control and injected embryos showing defect in development. Embryos injected with c*ldn-dΔYV* showed significant developmental defect compared to uninjected and embryos injected with c*ldn-d* and *cldn-a* (P-value: 0.0003). The percentage of c*ldn-dΔYV* injected embryos that showed developmental effect was (95.8± 5.89). n (*cldn-d:* 76, *cldn-dΔYV*: 169, *cldn-a:* 86, uninjected: 161). Error bars represent SD. ** : P-value ≤ 0.01, ** : P-value ≤ 0.01, *** : P-value ≤ 0.001, n.s.: non-significant. Scale bars: 50 µm.

### Electron microscopy showed TJ-like structures at early cleavage furrows

ZO1 protein has been also found outside of TJs *e.g.* in AJs (Junichi Ikenouchi, Kazuaki Umeda, Sachiko Tsukita, 2007; Yuji Yamazaki,*† Kazuaki Umeda,‡ Masami Wada,* Shigeyuki Nada & Shoichiro Tsukita, 2008). We therefore verified that the ZO1-and Buc-positive structures at the distal cleavage furrows of the 8-cell embryos form TJs using electron microscopy. Indeed, electron micrographs of 8-cell stage embryos showed electron-dense membrane sections resembling TJ-like structures at the cleavage furrows where germ plasm is localized (Fig. 7E). In contrast, we did not find these structures at the cleavage furrows, where germ plasm is not accumulated (Fig. 7F). This result shows for the first time that early zebrafish embryos have TJ-like structures already at the 8-cell stage, which are positive for ZO1 protein.

### The tight junction receptor Cldn-d anchors germ plasm

Claudins are one family of receptors, which physically connect the TJ in the epithelial and endothelial tissues of vertebrates. Claudins are transmembrane proteins that bind to the PDZ domains of scaffolding zonula occludens (ZO) proteins through their cytoplasmic C-terminal YV (Tyrosine-Valine) motifs (Furuse et al., 2014; Mccarthy et al., 2000). More than 50 claudins are identified in teleost fishes with restricted tissue expression patterns (Kolosov et al., 2013).

We therefore screened the list of potential BucLoc interactors for Claudin proteins. We detected Cldn-d as highly enriched in the pull-down assay (Supplementary table 1). In the Zfin database (https://zfin.org/) Cldn-d and -e are the only maternally expressed Claudins in zebrafish (Supplementary figure 10). Indeed, the *Xenopus* homolog Xcla is also maternally expressed (Brizuela et al., 2001), but its role in germ plasm tethering has never been investigated.

Based on the colocalization of Buc with TJs and the interaction of ZO1 with Cldn-d, we addressed the hypothesis that Cldn-d acts as a membrane anchor for germ plasm. We injected *cldn-d* mRNA into 1-cell zebrafish embryos expressing Buc-GFP from a transgene to allow the *in vivo* detection of germ plasm localization. Compared to uninjected control embryos the injection of *cldn-d* mRN*A* led to a significantly higher number of ectopic germ plasm spots at 2-3 hpf (Fig. 8A, B, E). As a specificity control, we injected the same concentration of mRNA encoding Cldn-a, but did not detect a change of germ plasm spots similar to uninjected controls (Fig. 8 A, C, E). The C-terminal amino acids Tyrosin and Valine are crucial for the interaction of Claudins with ZO proteins(Itoh et al., 2014). We therefore generated a Cldn-d mutant lacking the interaction motif (C-terminal YV, named Cldn-dΔYV), which was previously shown to act as a dominant-negative form of Claudin-receptors. Notably, *cldn-dΔYV* injected embryos showed a significantly reduced number of germ plasm spots in comparison to uninjected embryos (Fig. 8A, D, E). However, c*ldn-dΔYV* injected embryos displayed severe developmental defects in which the cells did not attach to each other (Fig. 8D, D’).

To exclude that the reduced number of germ plasm foci in *cldn-dΔYV* injected embryos is a secondary result caused by a defect in cell attachment, we targeted its expression to two blastomeres in a 16-cell embryo. Injection of *cldn-dΔYV* mRNA into 16-cell stage embryos still reduces the number of germ plasm spots. At this stage, the junctions are matured and the germ plasm containing cells can easily be distinguished from somatic cells, since they hold the central position in the marginal row of four blastomeres (‘middle blastomeres’). We injected *cldn-dΔYV* into two middle blastomeres surrounding one germ plasm spot (Fig. 9A) using uninjected and wild-type *cldn-d* injected embryos as controls. The number of Buc spots was counted right after injection (16-cell stage) and then followed up in regular time periods (see table 2 in the Supplementary). In this assay, embryos developed normally and did not show developmental defects (Fig. 9B, C). Interestingly, we still observed a significant reduction in the number of germ plasm spots in *cldn-dΔYV* injected embryos (Fig. 9B, C, D), compared to uninjected and *cldn-d* injected controls (Fig. 9D). More than 35% of *cldn-dΔYV* injected embryos lost a germ plasm spot. In contrast, only 6% of the embryos injected with *cldn-d* showed germ plasm spot reduction which could be accounted for the physical destruction during the injection process (Fig. 9D, table 1 in the Supplementary). Our results suggest that Cldn-d is involved in the recruitment of germ plasm to the TJs and the negative effect of the Cldn-dΔYV variant suggest that the C-terminal interaction domain is required to form the germ plasm complex at the early TJs at the cleavage furrow.

**Table 1:**
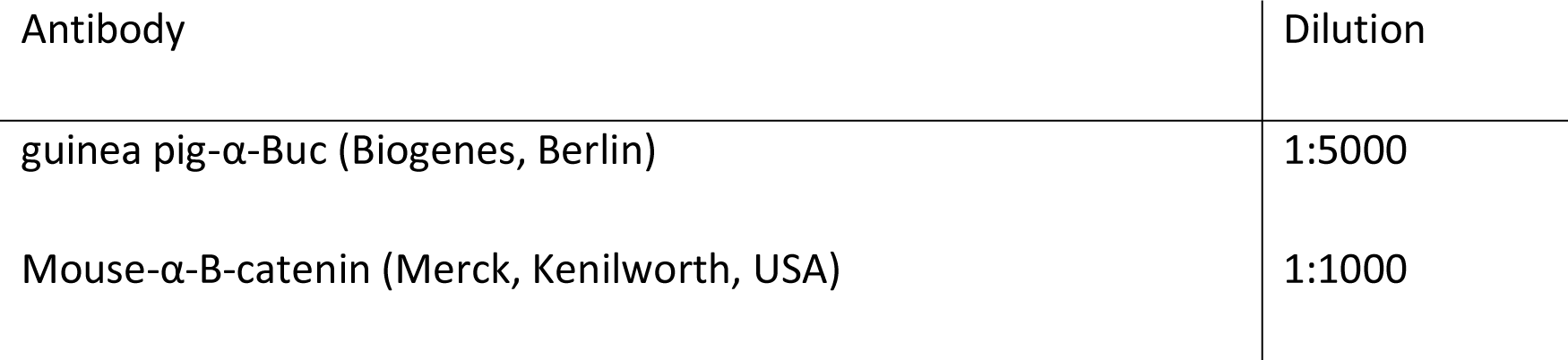
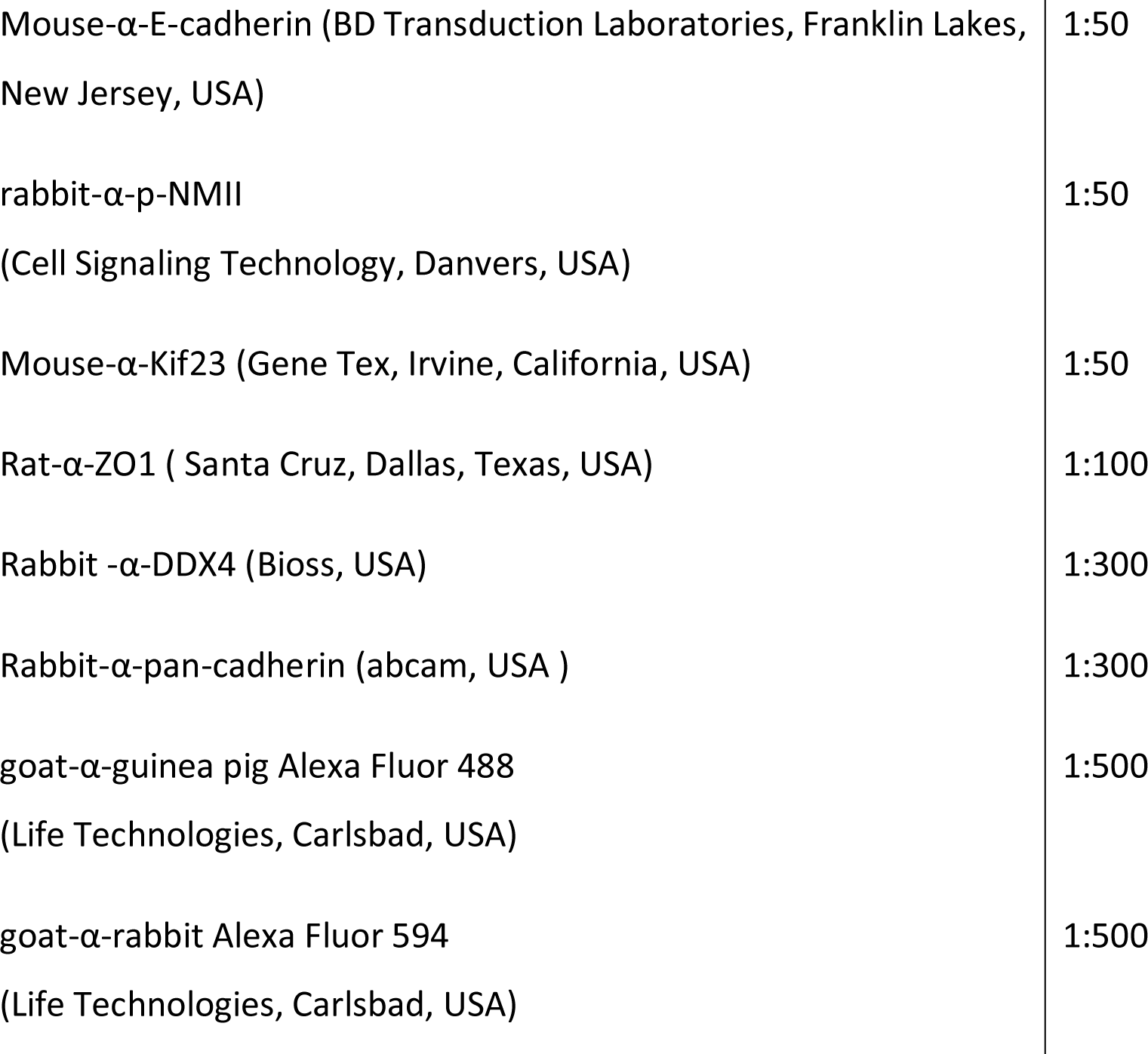
Antibodies used for immunostaining.

| Antibody | Dilution |
| --- | --- |
| guinea pig-α-Buc (Biogenes, Berlin) | 1:5000 |
| Mouse-α-B-catenin (Merck, Kenilworth, USA) | 1:1000 |
| Mouse-α-E-cadherin (BD Transduction Laboratories, Franklin Lakes, New Jersey, USA) | 1:50 |
| rabbit-α-p-NMII<br>(Cell Signaling Technology, Danvers, USA) | 1:50 |
| Mouse-α-Kif23 (Gene Tex, Irvine, California, USA) | 1:50 |
| Rat-α-ZO1 ( Santa Cruz, Dallas, Texas, USA) | 1:100 |
| Rabbit -α-DDX4 (Bioss, USA) | 1:300 |
| Rabbit-α-pan-cadherin (abcam, USA ) | 1:300 |
| goat-α-guinea pig Alexa Fluor 488<br>(Life Technologies, Carlsbad, USA) | 1:500 |
| goat-α-rabbit Alexa Fluor 594<br>(Life Technologies, Carlsbad, USA) | 1:500 |

**Table 2:**
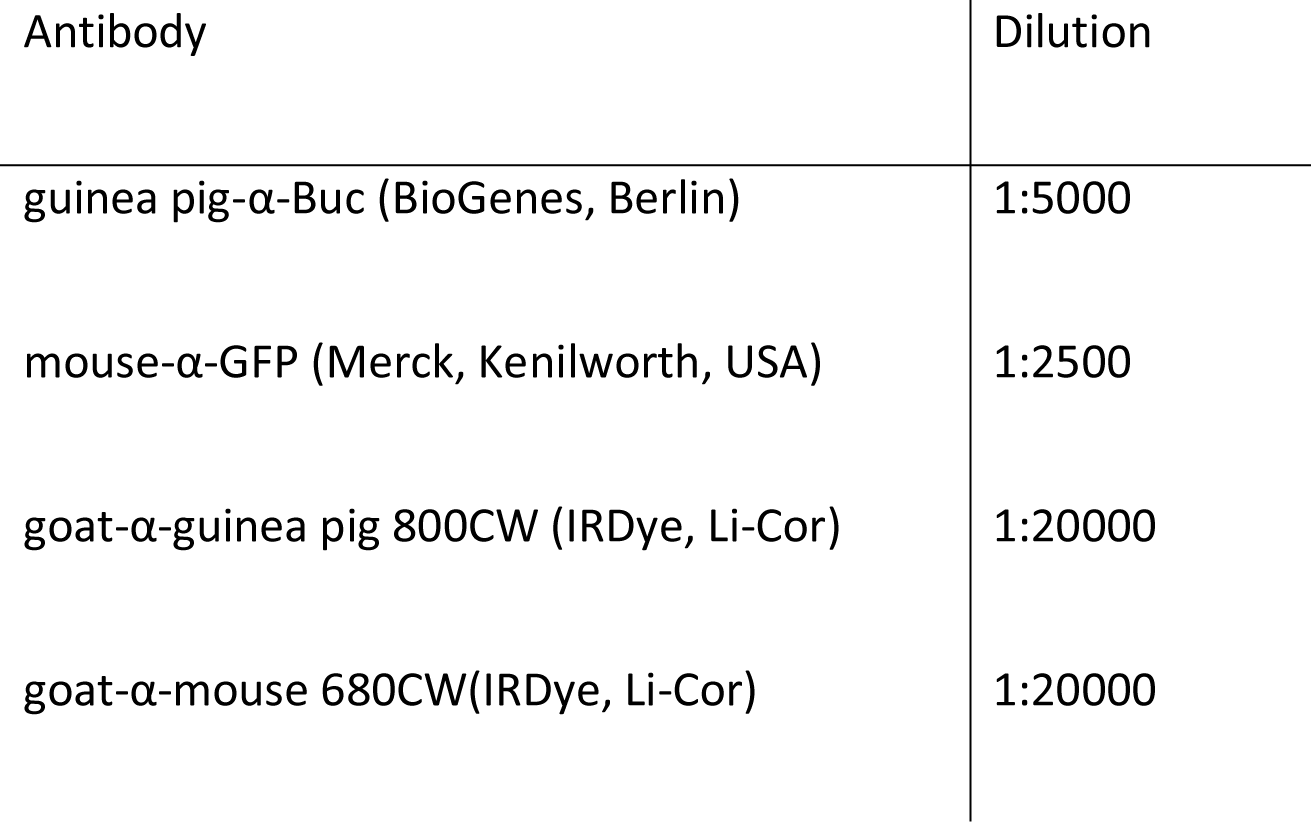
Antibodies used for western blotting.

| Antibody | Dilution |
| --- | --- |
| guinea pig-α-Buc (BioGenes, Berlin) | 1:5000 |
| mouse-α-GFP (Merck, Kenilworth, USA) | 1:2500 |
| goat-α-guinea pig 800CW (IRDye, Li-Cor) | 1:20000 |
| goat-α-mouse 680CW(IRDye, Li-Cor) | 1:20000 |

**Figure 9.**
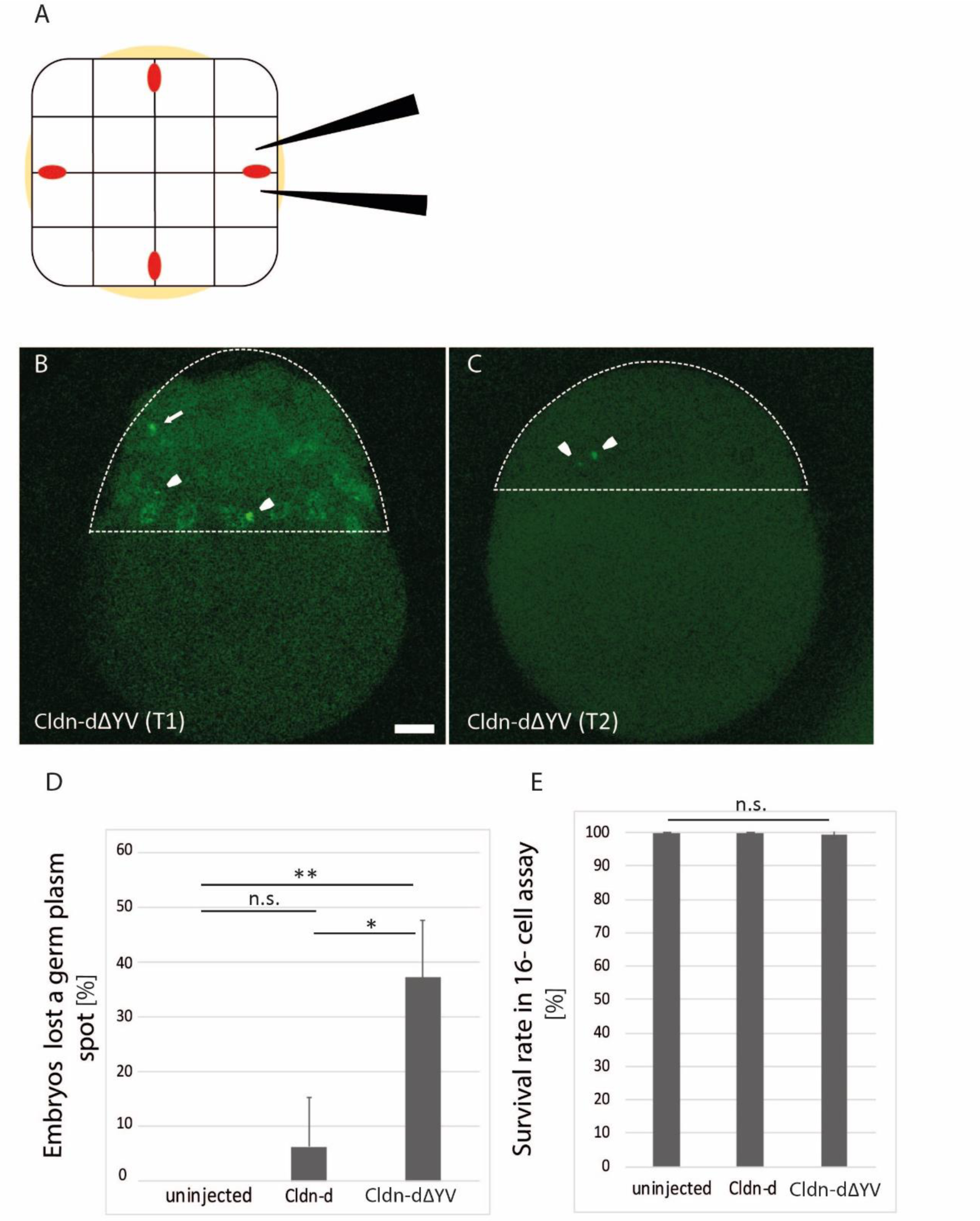
Cldn-dΔYV reduces the number of germ plasm spots. A) Schematic representation of 16-cell injection assay. The embryo is shown in animal view, germ plasm is shown as red spots. Two middle blastomeres surrounding a single germ plasm spot were injected. B) A *cldn-dΔYV* injected embryo from Buc-GFP transgenic line showing 3 Buc spots in lateral view. C) The same embryo shown in B at 2 hpf. One Buc spot disappeared (white arrow in B), while the other spots are sustained (arrowheads in B and C). D) Quantification of embryos which lost a germ plasm spot. c*ldn-dΔYV* injected embryos lost a germ plasm spot (35.1 ± 11.2%) which was significantly higher than c*ldn-d* injected (6.12 ± 10.8%; p=0.014) or uninjected (0 ± 0.0%; p=0,0024) embryos. No significant difference was seen between Cldn-d and uninjected embryos (P-value: 0.37). E) Percentage of embryo survival rate in 16-cell assay. There was no significant difference in the survival rate between injected and control embryos (P-value: 0.43). n (*cldn-d:* 49, *cldn-dΔYV*: 94, uninjected: (see supplementary table 2). Error bars represent SD. *** : P-value ≤ 0.01, ** : P-value ≤ 0.01, *, n.s.: non-significant. Scale bar: 50 µm.

Taking together, our results suggest that the germ plasm aggregates at localization foci of early formed TJs at cleavage furrows. The colocalization and protein-protein interaction data from the immunoprecipitations suggest that Buc together with other germ plasm components are linked to the early forming TJs via ZO1 protein and NMII recruitment using Cldn-d as membrane anchor (see model in Fig. 10).

**Figure 10.**
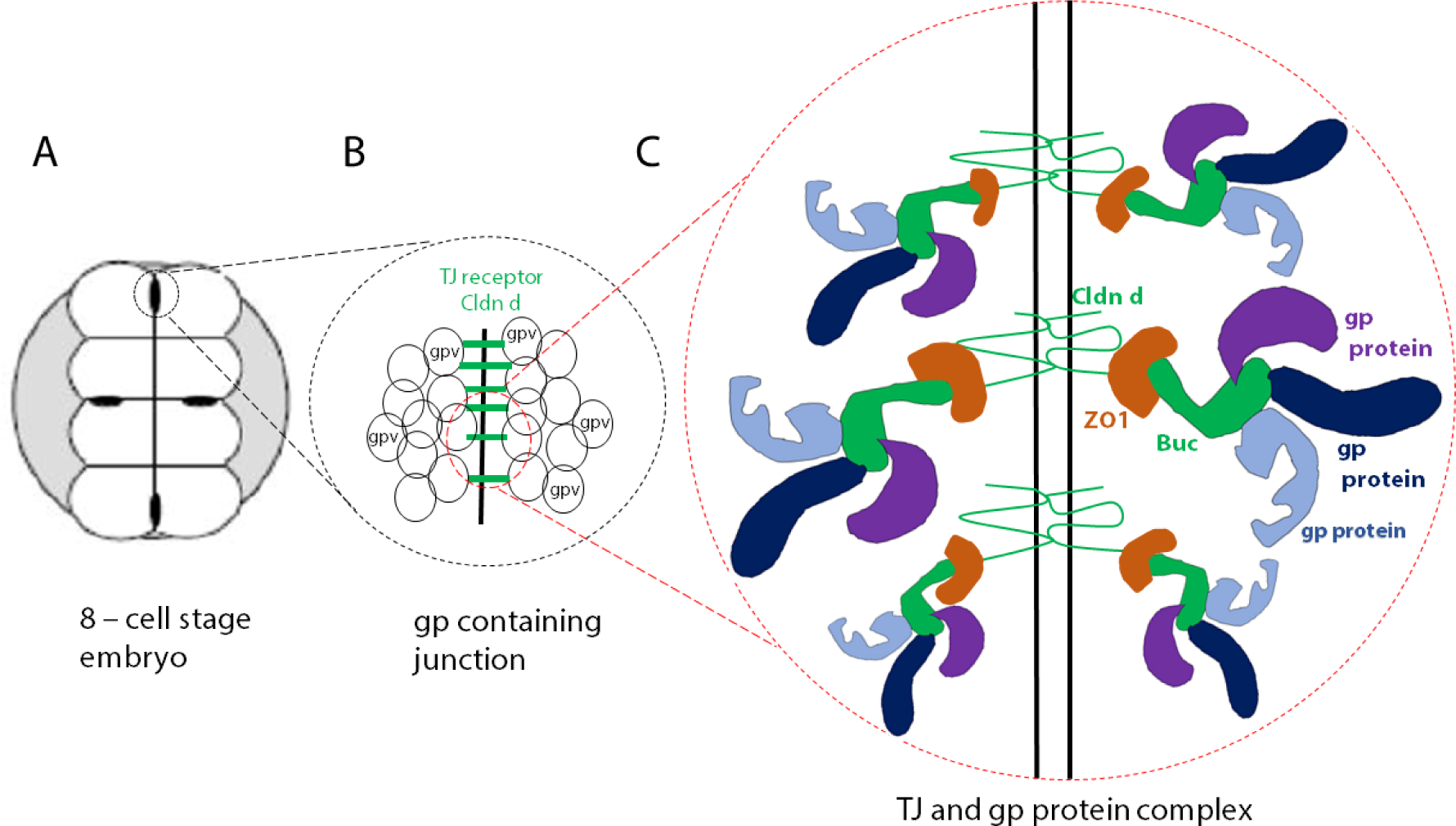
Proposed model for germ plasm localization in zebrafish. TJs anchor germ plasm at early cleavage furrows in zebrafish embryos. A) Schematic representation of a zebrafish embryo at 8-cell stage. Germ plasm spots are shown in four black spots. B) Magnification of a germ plasm spot from the embryo in (A). Vesicles containing germ plasm and TJ proteins (gpv) are anchored to the cleavage furrows by the TJ receptor Cldn-d (green). C) Representative magnification of the dashed red circle in (B). Buc in a complex with other germ plasm (gp) proteins interacts with the C-terminal end of Cldn-d. Note that it is currently not known whether Buc binds directly to ZO1.

## Discussion

Our data show that the localization machinery of germ plasm is different between vertebrates and invertebrates. We identified that the N-terminal Buc (aa11-88), is necessary and sufficient for the localization of the protein, the zebrafish germ plasm organizer. This region of Buc contains two PLDs, however they do not play a role in germ plasm localization in zebrafish. We found NMII as an interacting partner and colocalizing protein for Buc, suggesting that germ plasm might be anchored to one of the cellular structures through NMII. Indeed, we provide evidence that germ plasm localizes to TJs in early zebrafish embryos and overexpression of TJ receptor protein Claudin-d induces ectopic germ plasm formation.

### Evolutionary conservation of germ plasm anchorage among vertebrates

Invention of multicellularity requires cell adhesion and germ plasm for sexual reproduction. With our finding, it will be possible to address whether germ plasm localization at TJs was already used at the origin of Metazoa or whether it is a derived mechanism acquired during vertebrate evolution. The isolated BucLoc motif does not show homologies to known protein domains, which did allow to deduce its biochemical function. However, it proposes vertebrate species, which might use a similar localization mechanism for Buc like the zebrafish. Indeed, we found with this approach that the germ plasm in the early chicken embryo also localizes to the cleavage furrow. As the BUVE-motif containing BucLoc shows similarity to Buc-like ORFs in the genome of Laurasiatheria mammals (Bontems et al., 2009; Krishnakumar et al., 2018), it would therefore be exciting to study the localization of TJ proteins in their embryos during early cleavages.

Our data suggest that the tethering of germ plasm is conserved among vertebrates, as we observed identical positioning in *Xenopus,* zebrafish and chicken, but not in *Drosophila*, suggesting a similar anchorage mechanism in fish, frogs and chicken. In contrast, *Drosophila* sOsk is not targeted by the vertebrate localization system. Therefore, our results show that the localization machinery of germ plasm is different between vertebrates and invertebrates. Despite the functional equivalence of Buc and Osk which is previously shown (Krishnakumar et al., 2018), we here provide evidence that Buc and Osk proteins use different mechanisms to localize germ plasm. Germ cell specification activity of these germ plasm nucleators looks conserved, whereas the mechanism of their localization seems to adapt to the architecture of the embryo. Therefore, different localization mechanisms are consistent with the different shapes of early embryos.

### Germ plasm localizes at TJs at early cleavage furrows in zebrafish embryos

Furrow-associated microtubule-array (FMA) is previously described as cytoskeletal structure tethering germ plasm in zebrafish (Jesuthasan, 1998; Pelegri et al., 1999). However, the exact cellular structure responsible for the anchorage of the germ plasm in zebrafish was still unknown. It was previously shown that NMII is involved in various cellular structures (Liu et al., 2012; Nair et al., 2013; Vicente-Manzanares et al., 2009). NMII is activated via phosphorylation by a number of protein kinases such as Rho associated protein kinase (ROCK). This ultimately results in the assembly of myosin filaments, which is also shown to colocalize with germ plasm RNA during zebrafish embryonic development (Amano et al., 1996; Miranda-Rodríguez et al., 2017; Nair et al., 2013). Our results showed that (i) myosin light chain co-immunoprecipitated with Buc and that p-NMII is colocalizing with Buc protein, suggesting that germ plasm might get anchored to one of the cellular structures through NMII (Fig. 6). (ii) Germ plasm colocalizes with the TJ protein ZO1, which is a known direct binding partner of Claudins at the TJs(Itoh et al., 2014) (Fig. 7). (iii) Co-immunoprecipitations of Buc co-purified ZO and Claudin (Fig. 6). (iv) EM-micrographs show the presence of TJ-like structures at the cleavage furrows in the 8-cell zebrafish embryo (Fig. 7). (v) *cldn-d* injection causes the formation of a higher number of germ plasm spots, whereas Cldn-d with a mutated interaction motif for ZO proteins (C-terminal YV motif) functions as a potential dominant negative resulting in fewer germ plasm spots (Fig. 8; Fig. 9). These results strongly support the model that newly forming TJs at the cleavage furrows represent the anchorage hub for the germ plasm in zebrafish. This localization might be critical to achieve a threshold concentration for phase-transition, which plays an important role during germ plasm aggregation (Kistler et al., 2018; Krishnakumar et al., 2018; Trcek & Lehmann, 2019).

### TJs as an anchorage hub for germ plasm

Anchoring of the germ plasm to the TJs is different from anchoring to the posterior cell cortex in *Drosophila* oocytes and embryos. The posterior localization of germ plasm is essential for the specific embedding into budding PGC at the posterior pole in early *Drosophila* embryos. However, in the zebrafish embryos the role of germ plasm anchorage is different. The observed accumulation and linkage to the TJs keep the germ granules concentrated in one spot of the cytoplasm, thereby inhibiting symmetric distribution of germ plasm during the following cell divisions. This anchorage results in the preservation of only four PGC up to the 512 cell stage. Only after the mid-blastula transition (MBT) at cell cycle 10, germ plasm is symmetrically inherited when the PGC start to divide forming four clusters of PGCs (Dosch, 2015; Knaut et al., 2000; Wolke et al., 2002). At that time the germ plasm is localized into perinuclear clusters, enabling a symmetric distribution during PGC divisions (Strasser et al., 2008). Therefore, the anchorage to the TJs has to be released at MBT, as the PGC start dividing and the germ plasm needs to be inherited symmetrically by both daughter cells to ensure their fate.

A release of the germ plasm aggregate from the TJ could be a consequence of the modification of Claudins, Buc or of other unknown bridging protein(s). We believe that our results show a specific function of Cldn-d, as injection of Cldn-d caused a significant increase in the number of germ plasm spots, whereas Cldn-a had no effect. Furthermore, co-immunoprecipitation experiments revealed a specific interaction of Buc with Cldn-d but not with Cldn-a. This differential biological activity could be caused by the particular capability of Cldn-d to form *de novo* TJs in the early cleavage furrows, but we would rather favor a model in which the interaction of Cldn-d with the germ plasm is specific. This model could explain the release of the germ plasm from TJs by exchange or dilution of the maternal Cldn-d with other claudins, starting at the onset of zygotic expression. Future experiments have to address the mechanism by which germ plasm is inherited symmetrically into both daughter PGCs after MBT.

### Function of Buc in germ plasm anchorage at the TJs

Sequence analysis of Buc did not reveal any characterized domain within the protein (Bontems et al., 2009; Krishnakumar et al., 2018). However, sequencing of *buc^p106re^* allele revealed a mutation in the 6th exon of Buc genomic locus, which would cause a deletion of only 37 C- terminal amino acids, suggesting an essential role of its C-terminal end. Within this C-terminus Arginine residues are dimethylated, thereby enabling the direct interaction of Buc with the zebrafish Tudor homologue Tdrd6 (Roovers et al., 2018). Tdrd6 interacts with the known RNAs enriched in the germ plasm and it was shown that it is involved in the loading of germ plasm components into PGCs. Furthermore, high-resolution microscopy showed that Tdrd6 and Buc form particles with germ plasm mRNA in which Buc is localized in the core of the particle, whereas Tdrd6 is mostly positioned at the periphery of the particles at the 4-cell stage when the germ plasm start to accumulate at the cleavage furrows. These data suggested that Buc cooperate with Tdrd6 like Osk with Tudor in the aggregation of the germ plasm.

Buc is also interacting with Vasa as shown by co-immunoprecipitations and experiments using split Cherry, suggesting that Buc and Vasa bind directly within the N-terminal 360 amino acids (Krishnakumar et al., 2018). Furthermore, the N-terminal half of Bucky Ball is also able to interact with *nanos3*-3’-UTR RNA. However, it is unknown whether Buc can directly interact with mRNA of the germ plasm, because it does not contain any characterized RNA binding motif.

Sequence comparison with 15 related Buc proteins revealed a conserved 100 amino acid N-terminus, which was named BUVE motif (Buc-Velo) (Bontems et al., 2009). The BUVE domain was shown to be essential for the formation of the amyloid-like aggregates in the BB in Xenopus oocytes. The BUVE domain contain potential prion like domains (PLD) (Alberti et al., 2009), which were shown to be essential for the aggregation process, based on the fact that the replacement of critical residues with charged amino acids inhibited the aggregation. These results suggested that the BUVE domain of Velo1 and Buc is required for amyloid-like germ plasm aggregation in the BB. However, Velo1 variants in which the potential PLDs were replaced with unrelated PLDs were inactive, whereas the replacement with the related sequences from zebrafish Buc were active, revealing a sequence specificity (Boke et al., 2016). Surprisingly intrinsic disorder prediction of Buc showed that N-terminus (aa 1–150) is the largest ordered sequence in Buc (Krishnakumar et al., 2018). *Drosophila* Osk was recently shown to form aggregates in an ectopic system (insect S2 cells) although Osk does not display any PLD domains (Boke et al., 2016; Jeske et al., 2017). Nevertheless, we tested for the role of the predicted PLDs within the N-terminus when we identified the region between aa11 and 88 to be essential and sufficient for the localization of Buc to the 4 germ plasm spots at the cleavage furrows (Fig. 4). The detailed mapping showed however, that none of these two potential PLDs within this sequence are essential, but short stretches C-terminal of them (Fig. 5). This result does not exclude that the PLDs are involved, as the additional regions might be required for the proper presentation of the PLDs. However, co-immunoprecipitation and Mass spectrometry analyses showed that the domain between aa 11-88 interacts with about 213 peptides including myosin light chain and Cldn-d (Fig. 6). Nonetheless, 213 peptides is a high number of interactions and includes probably a number of unspecific interactions. For instance, we purified five of ten subunits of the RNA-exosome complex and probably only one subunit directly interacts with Buc. We therefore prefer to interpret these results in a different way, whereby the BUVE domain including the BucLoc represents a protein-protein interaction module that indirectly enables the formation of a protein RNA complex essential for germ plasm aggregation. Future experiments will identify the mechanisms and the direct interaction partners of Buc.

### Buc and TJs in biomolecular condensates

Increasing evidence suggest that germ granules in many different organisms are formed by phase separation. Germ plasm consists of spherical units of protein RNA aggregates that show a highly dynamic exchange with the surrounding cytoplasm (recently reviewed in (Dodson & Kennedy, 2020; So et al., 2021)). Indeed, the BucLoc motif was previously shown to play a crucial role in aggregating the Balbiani body in the Xenopus oocyte, which is probably the largest biomolecular condensate in the animal kingdom (Boke et al., 2016). However, our results show that the Prion-like domains in the BucLoc motif, which control Balbiani body assembly, are not required for germ plasm anchoring in the embryo. Nonetheless, our previous data show that Buc forms liquid-like condensates in the embryo (Krishnakumar et al., 2018; Riemer et al., 2015; Roovers et al., 2018),suggesting that aggregation does not control its embryonic localization.

Interestingly, ZO Proteins also induce the assembly of liquid-like condensates (reviewed in (Canever et al., 2020; Citi, 2020)). The condensation of ZO proteins in cell culture and zebrafish embryos induces the assembly of TJs revealing an unexpected activity in the cytoplasm to control the formation of TJs (Beutel et al., 2019; Schwayer et al., 2019). Our finding that Buc and ZO1 colocalize, raises the question, whether Buc indeed autonomously induces condensates or whether this activity is mediated by ZO1. However, we previously showed that Buc condensates in HEK293 cells, which do not form TJs supporting Buc’s autonomous phase separation activity (Krishnakumar et al., 2018).

Fascinatingly, we show that the injection of Cldn-d had a similar activity on forming extra germ plasm spots compared to the injection of Buc (Bontems et al., 2009). This effect is unexpected considering the instructive role of ZO-protein on TJ formation (Fig. 8A, B). Two scenarios could explain this strong activity of Cldn-d. Either Cldn-d induces the expression of Buc-GFP or it inhibits it degradation. As there is no transcription between the injection of Cldn-d mRNA and the 2 hpf, we do not favor increased expression, although we cannot exclude it. Rather we propose that Buc anchored at TJs is stabilized, while unlocalized Buc-GFP granules are degraded. This is consistent with previous time-lapse experiments, in which unlocalized Buc-GFP granules are cleared around the 32-64 cell stage (Riemer et al., 2015). In this scenario, the maternal load of Cldn-d is limited and just sufficient to form four spots. Indeed, the loss of spots after injection of dominant-negative Cldn-d seems to support this hypothesis. Taken together, our results support an essential function of Buc generating a multiprotein hub at the newly forming TJs that allows the anchorage of germ granules.

### Conclusion

In conclusion, we found that vertebrates and invertebrates utilize different germ plasm localization mechanisms, with evolutionary conservation between vertebrates. We discovered that TJs anchor germ plasm during early zebrafish embryogenesis and that germ plasm in zebrafish is anchored to the TJs and via Cldn-d receptor protein. Microinjection of *cldn-d* induced extra germ plasm spots. Currently we believe that NMII interaction with Buc drives the localization of germ plasm into the TJs, as shown in the following model (Fig. 10).

## Materials and methods

### Zebrafish handling and manipulation

Zebrafish (*Danio rerio*) was used as an animal organism in this study, AB*TLF (wild-type) and Buc-GFP transgenic zebrafish line(Riemer et al., 2015). Fish were raised and maintained according to the guidelines from (Westerfield, 2000) (Westerfield M., 2000) and regulations from Georg-August University Goettingen, Germany.

### Microinjection

Previously synthesized capped RNA was diluted with 0.1M KCl and 0,05% phenol red (Sigma Aldrich, Hannover). 2nl of RNA was injected into 1-cell stage embryos using PV820; WPI injecting apparatus (Sarasota, USA). Injected embryos were incubated in E3 medium at 28°C until they reached the developmental stage of the phenotype evaluation.

### 16-cell injection assay of Cldnd-ΔYV

To study whether non-functional Cldn-d has an influence on matured TJs, we conducted *cldn-dΔYV* injections in 16-cell embryos. In this assay, we injected the RNA directly into two cells next to a germplasm localizing tight junction. As a control we used uninjected and *cldn-d* RNA injected embryos. The number of Buc spots was counted right after injection and then followed up in regular time periods. Detailed description of the injection procedure at 16-cell stage is previously published (Bontems et al., 2009; Krishnakumar et al., 2018).

### *Drosophila* handling and manipulation

Flies were kept and crossed at room temperature or 25 °C. To collect embryos, the flies were kept in cages with apple juice agar plates at 25 °C. Experiments were approved by the Lower Saxony State Office for Consumer Protection and Food Safety (AZ14/1681). The pUASp *bcd*3’UTR plasmid expressing sOsk (Tanaka & Nakamura, 2008) was used to replace the *sosk* ORF with Buc ORF-GFP. A germline-specific *mat -Gal4VP16* driver was used to express UASp-based transgenes in oogenesis. Antibody staining and fluorescent *in-situ* hybridization was performed as described (Pflanz et al., 2015). The antibodies used were anti-PY20 (1/500, Biomol), rabbit anti-GFP (1/1000, Synaptic Systems, Göttingen, Germany) and anti-Vasa (1/5000 (Pflanz et al., 2015)). Anti-mouse and anti-rabbit antibodies coupled to Alexa 488, 568 or 647 were used as secondary antibodies (Invitrogen, 1/1000). Embryos were imbedded in DPX to provide clearing and to protect from bleaching.

### Biochemical methods

#### Co-immunoprecipitation (Co-IP)

CO-IP was performed to identify Buc protein interactome. Each sample was prepared from 500 deyolked high stage embryos after homoginization on ice in lysis buffer (10 mM Tris (pH 7.5), 150 mM NaCl, 0.5 mM EDTA, 0.5 % NP-40, 1x complete protease inhibitor cocktail (Roche, Mannheim)). The supernatant was subsequently used for the Co-IP using a GFP-binding protein coupled to magnetic beads (GFP-Trap_M; ChromoTek, Planegg-Martinsried) following manufacturers instructions. After pulling down, the magnetic beads and their bound proteins were either incubated with 2x SDS loading buffer for 5 min at 96 °C and analyzed via SDS-PAGE and western blotting or sent for mass spectrometry (Core Facility of Proteome Analysis, UMG, Goettingen), as described previously (Krishnakumar et al., 2018).

#### Selection criteria for specifically interacting proteins

In total, 3464 protein candidates interacted. From those, 1817 candidates were identified that interacted with both Buc-GFP and BucLoc-GFP. We were not interested in every candidate for interaction with Buc-GFP, as they might interact with any other region outside of BucLoc. Therefore, we applied a set of criteria to identify significant interacting candidates with BucLoc. First, any peptide below a background threshold of five in BucLoc-GFP was considered as not significant and were sorted out. Furthermore, only proteins with counts in BucLoc-GFP that were at least twice as high as in the negative control GFP were considered as significant. To further reduce overexpression artefacts, enrichment in the positive control and in the sample had to be within a magnitude of +/-4-fold. Applying these selection criteria, the number of potential BucLoc interaction proteins could be restricted to 213 interaction candidates (see the Supplementary for the full list of mass spectrometry candidates).

#### Immunohistochemistry

Embryos were fixed and stained as previously described (Riemer et al., 2015) with the following antibody concentrations.

#### Imaging

Images were taken by SteREO Lumar.V12 (Carl Zeiss Microscopy, Göttingen) and LSM780 confocal microscope and analyzed with xio Vision Rel. 4.8 software and ZEN2011 software (Carl Zeiss Microscopy, Göttingen), as described before (Riemer et al., 2015). Electron microscopy was performed at the facility for transmission electron microscopy (Max Planck Institute for Biophysical Chemistry, Göttingen).

#### Western blotting

Western blotting was performed to detect the specificity of Buc antibody as described before (Krishnakumar et al., 2018). Fluorescent signal was detected with Li-Cor Odyssey CLx Infrared Imaging system (Li-Cor, Lincoln, USA) and analyzed with the Image Studio Software (Li-Cor, Lincoln, USA).

#### In-vitro translation

Proteins were synthesized with the TnT SP6 Quick Coupled Transcription/Translation System (Promega, Madison, Wisconsin, USA).

#### Molecular biology methods Cloning

The template of all the constructs that are used in this study were amplified from reverse transcribed cDNA which was made from total ovarian RNA. Constructs are cloned with either restriction digestion or gate way cloning.

#### Bioinformatics Sequence alignment

Pairwise sequence alignment was used to compare protein sequences, using Needleman-Wunsch algorithm with the EMBL-EBI alignment software EMBOSS Needle (McWilliam et al., 2013).

#### PLD prediction

Fold amyloid (Fernandez-Escamilla et al., 2004), APPNN (Família et al., 2015), FISH amyloid (Gasior & Kotulska, 2014), and Aggrescan (Conchillo-Solé et al., 2007) algorithms were used to predict PLDs in BucLoc.

#### Analysis of mass spectrometry data

Overlaps in protein interactions between each Co-IP sample were analyzed using a Venn diagram generator (http://jura.wi.mit.edu/bioc/tools/venn3way/index.php). The Kyoto

Encyclopedia of Genes and Genomes (KEGG, http://www.genome.jp/kegg/) has been used to classify the BucLoc-GFP interaction candidates in collaboration with Dr. Thomas Lingner.

### Statistics

All the statistical analysis of the experiments have been carried out in Microsoft Excel and the Prism software (GraphPad Software, La Jolla, USA). Error bars indicate the standard deviation of averages. For each injection experiment, at least three independent replicates were used.

## Acknowledgements

We are thankful to Prof. E. A. Wimmer for providing the facilities to perform this research and G. Kracht for technical assistance.

## Supplementary data

**Supplementary figure 1.**
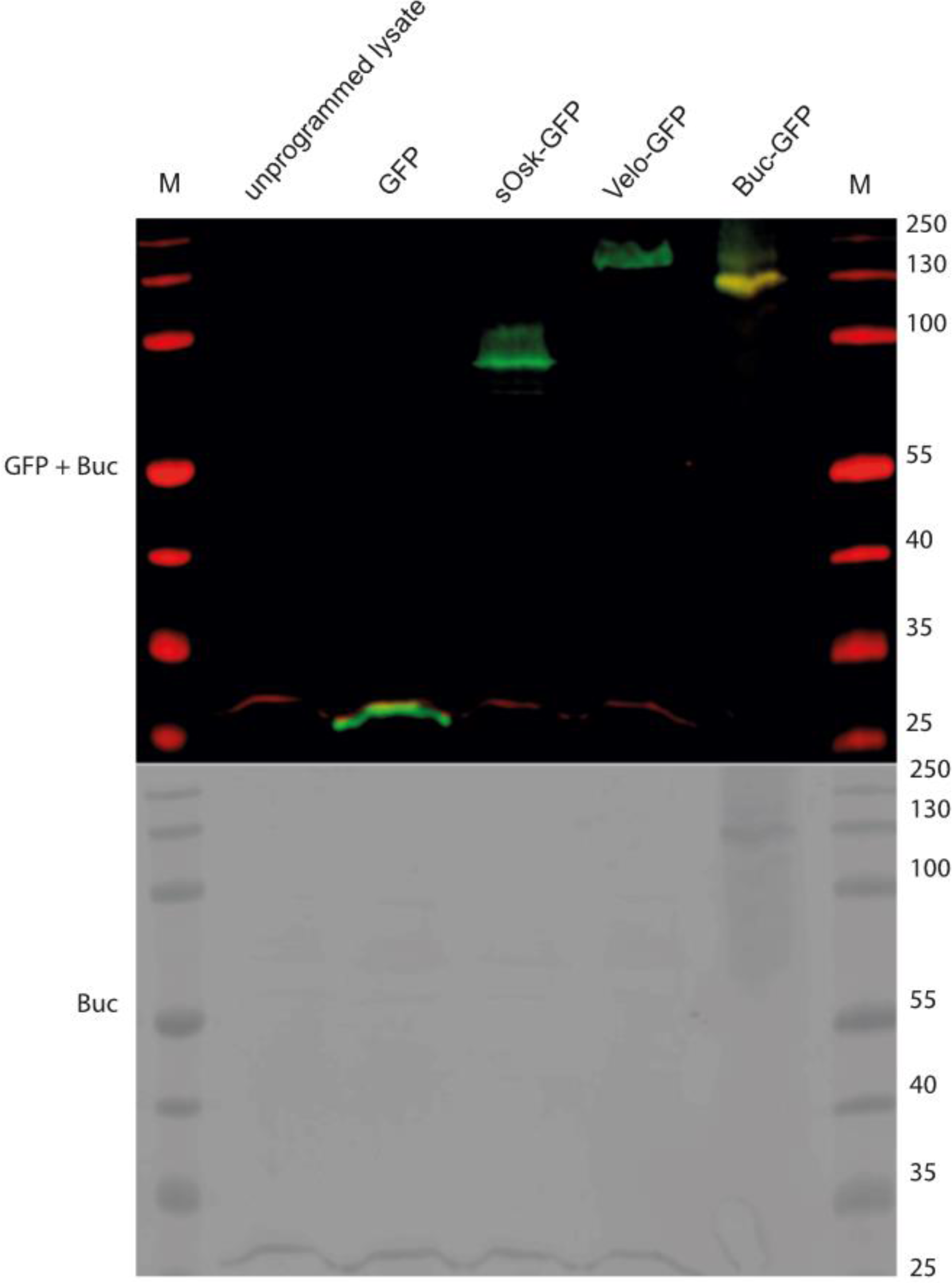
The Buc antibody does not cross-react with GFP, *Xenopus* Velo or *Drosophila* Oskar. Western blot showing anti-Buc (red in upper panel and black in lower panel) or anti-GFP (green in upper panel) antibody staining of *in vitro* translated GFP, sOsk-GFP, Velo-GFP and Buc-GFP. Unprogrammed lysate was used as negative control for protein translation. Buc-GFP is visualized by both anti-Buc and anti-GFP antibodies (yellow in merged panel and black in lower panel), whereas Velo-GFP, sOsk-GFP and GFP are only recognized by anti-GFP antibody, but not by anti-Buc antibody.

**Supplementary figure 2.**
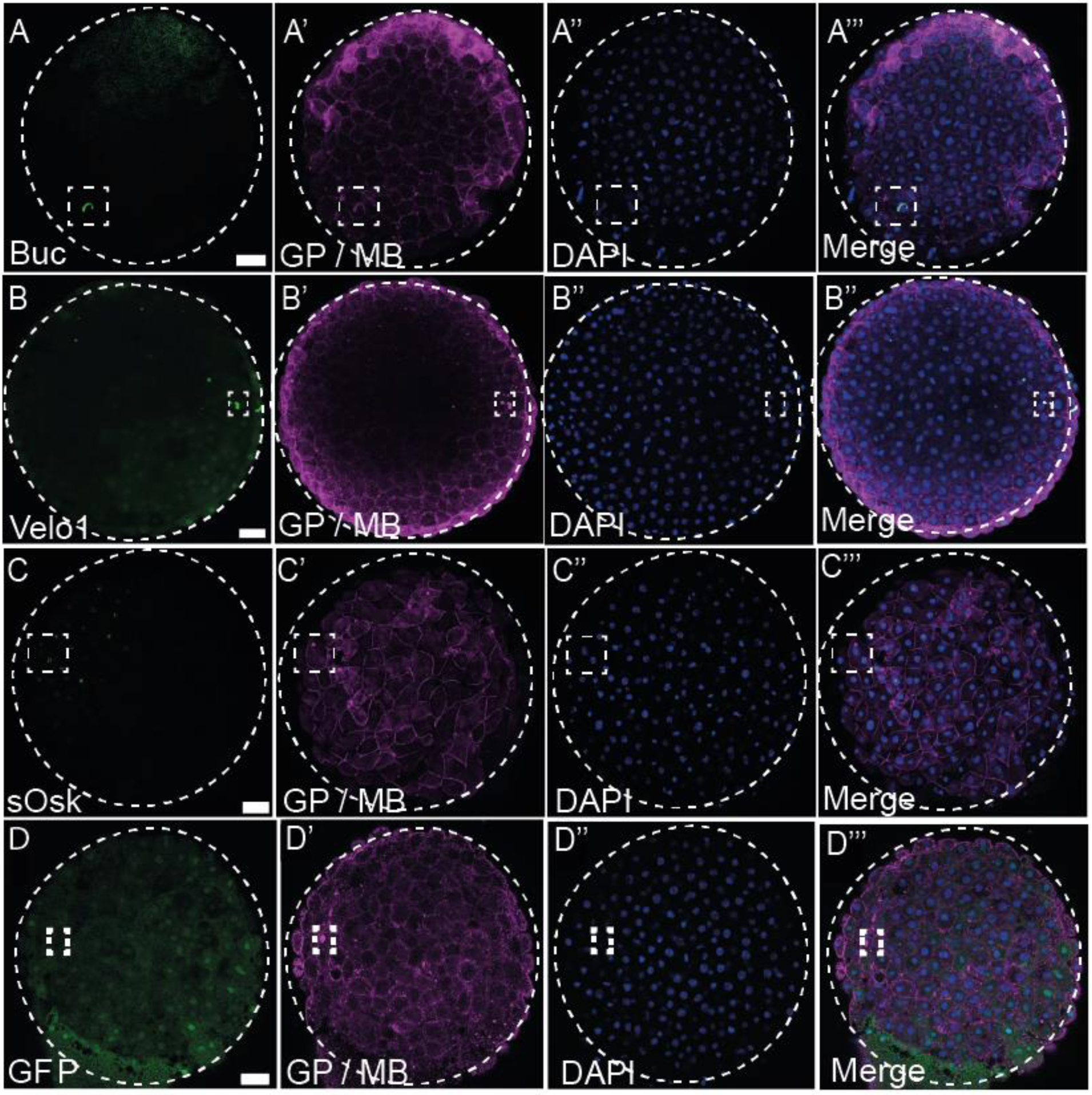
Germ plasm localization is conserved in vertebrates. Panels (A,B,C,D) show embryos at high stage from the animal view. Dotted circles outline the embryos. Dotted rectangles show magnified areas in figure 1. Colocalization of the GFP with endogenous Buc was determined by immunohistochemistry: 1st column – injected GFP fusions (green), 2nd column – endogenous Buc and beta-catenin (magenta), 3rd column – DAPI (blue) and 4th column – merge. Buc-GFP (A-A’’’) and *Xenopus* Velo1 (B-B’’’) colocalize with endogenous germ plasm, whereas *Drosophila* Osk(C-C’’’) shows nuclear localization. The GFP control shows ubiquitous low level fluorescence (D-D’’’). Scalebars: 50 µm.

**Supplementary figure 3.**
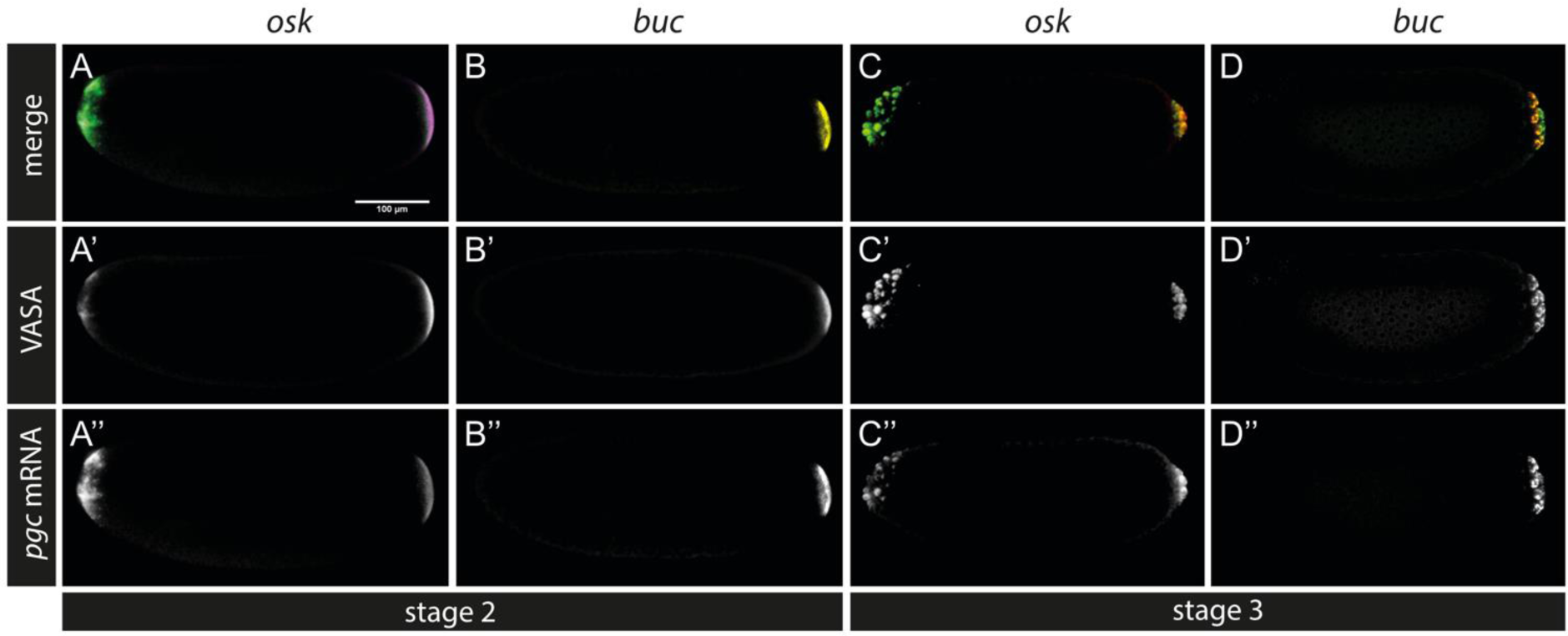
Vasa protein and pgc mRNA recruitment in transgenic flies. Vasa protein and *pgc* mRNA labeling of the transgenic embryos showed that sOsk specified ectopic PGCs, whereas Buc transgenics did not recruit Vasa protein or *pgc* mRNA at the anterior pole. (A-A’’, B-B’’) show VASA and *pgc* labeling of sOsk and Buc in stage2 transgenic flies. (C-C’’, D-D’’) show VASA and *pgc* labeling of sOsk and Buc in stage3 transgenic flies. Scale bar: 100 µm.

**Supplementary figure 4.**
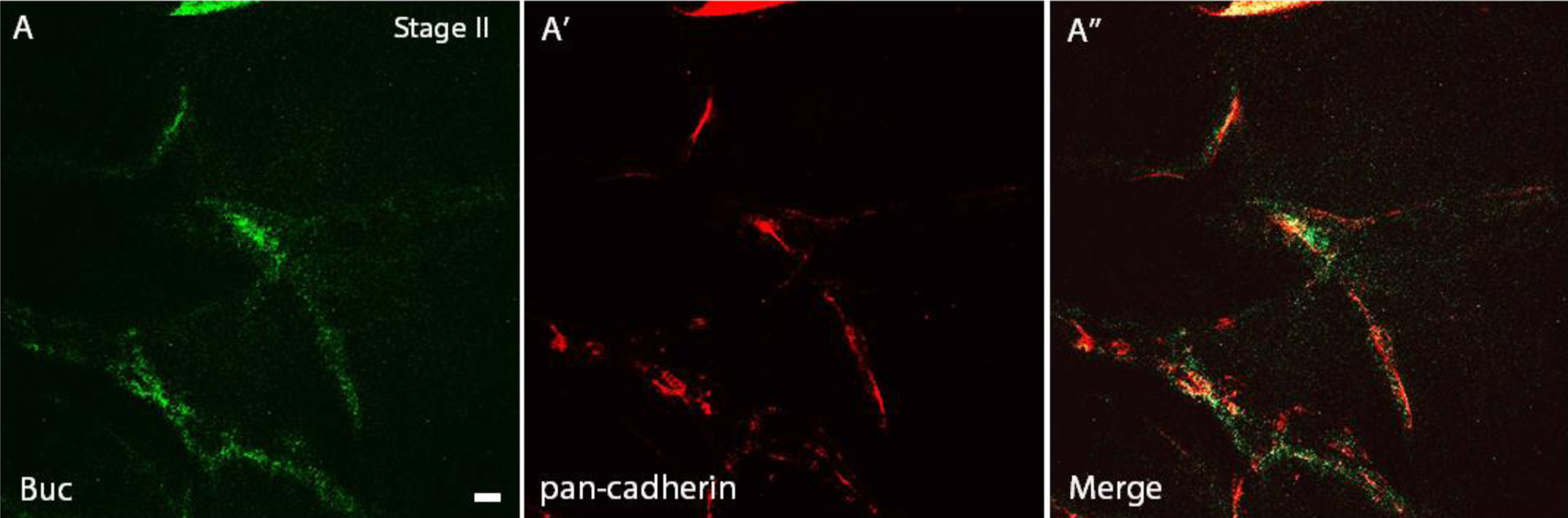
Buc localizes to the cell membrane in amniotes. Colocalization of Bucky ball (Buc) with cell membrane specific marker pan-cadherin was determined by immunostaining in stage II chicken embryos (4 hpf). Note the colocalization at cleavage furrows in stage II chicken embryos (4 hpf). (A) Buc (green), (B’)p-cadherin (red), (A’’) merge (yellow). Buc and p-cadherin are not located in the same protein granules as it was recorded for Buc and Cvh colocalization. Scalebar: 20 µm.

**Supplementary figure 5.**
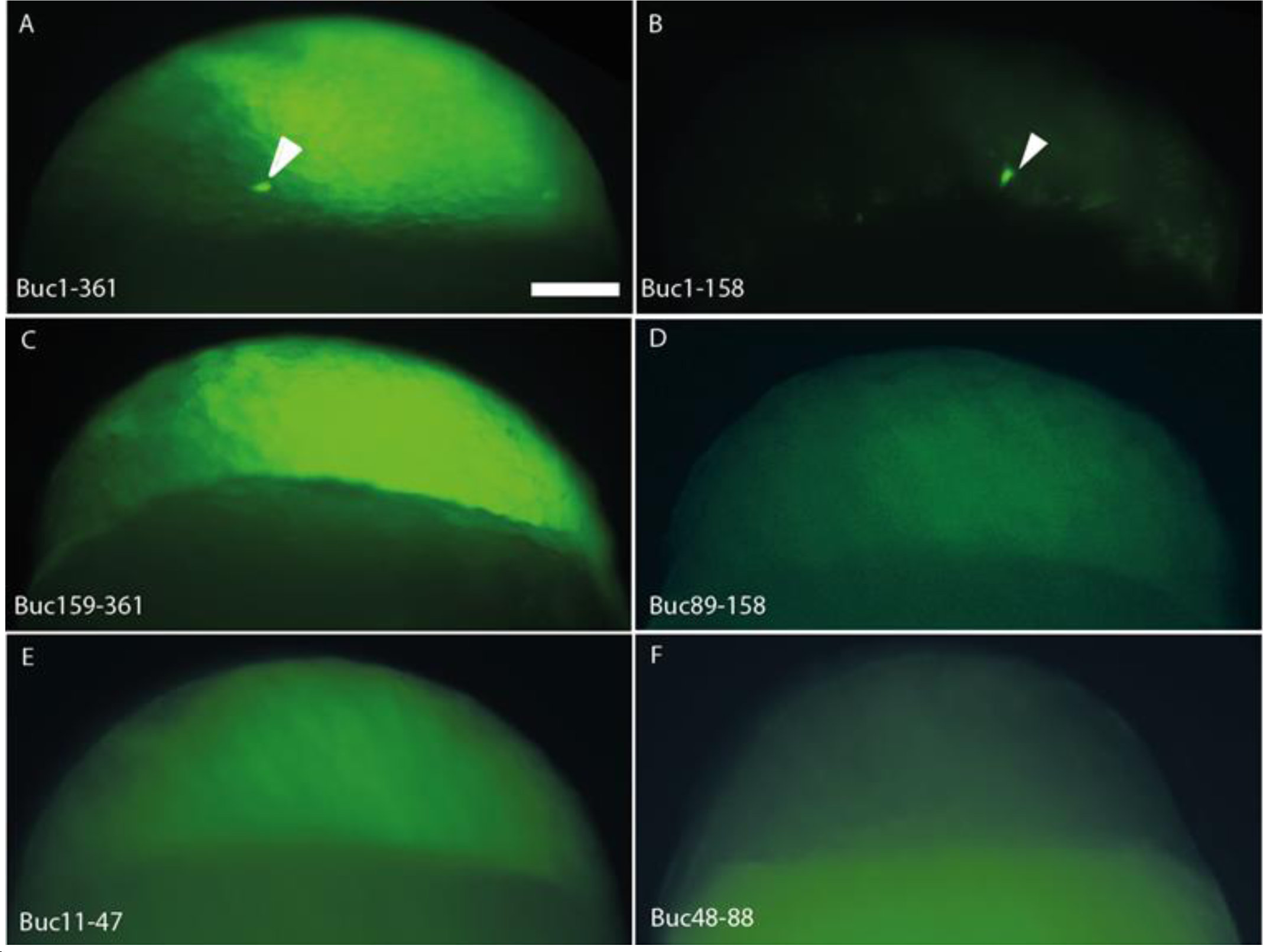
Systematic mapping of Buc localization motif. (A) N-terminal fragment of Buc (aa1-361) (white arrowhead) (100%). (B) Buc1-158 localizes (white arrowhead) (100%). (C) Buc159-361 is ubiquitous (0%). (D) Buc89-158 is ubiquitous (0%). (E) Buc11-47 is ubiquitous (0%). (F) Buc48-88 is ubiquitous (6.0±6.7%). Scale bar: 50 µm

**Supplementary figure 6.**
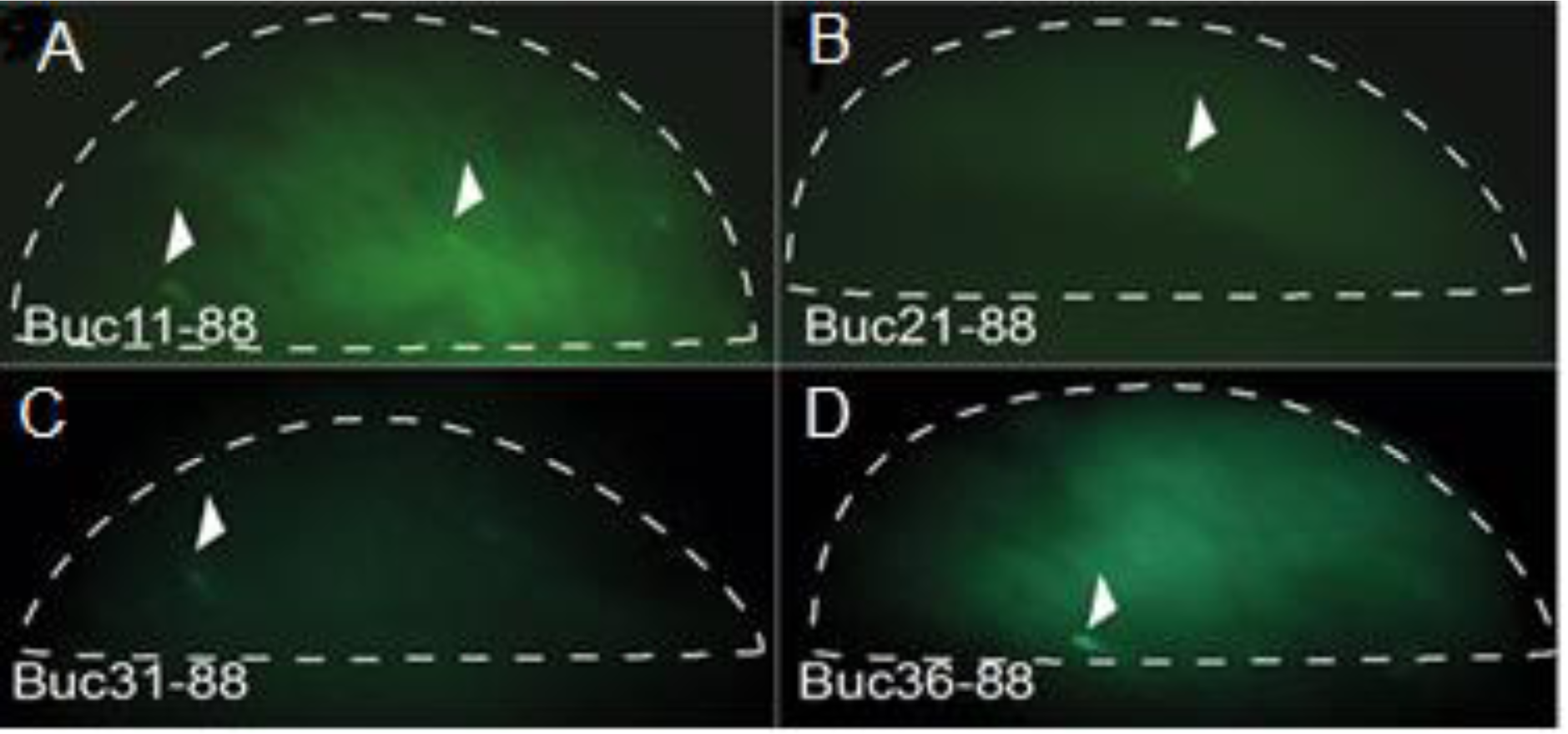
Aggregation and localization of BucLoc are separate activities. (A-D) Show embryos at high stage from the lateral view. Embryos are outlined by the dashed white line. Injected constructs showed fluorescent aggregates (white arrowheads). (A) Buc11-88 (91.4±6.8%). (B) Buc21-88 (67±4.0%). (C) Buc31-88 (60.1±7.9%). (D) Buc36-88 (52.2±13).

**Supplementary figure 7.**
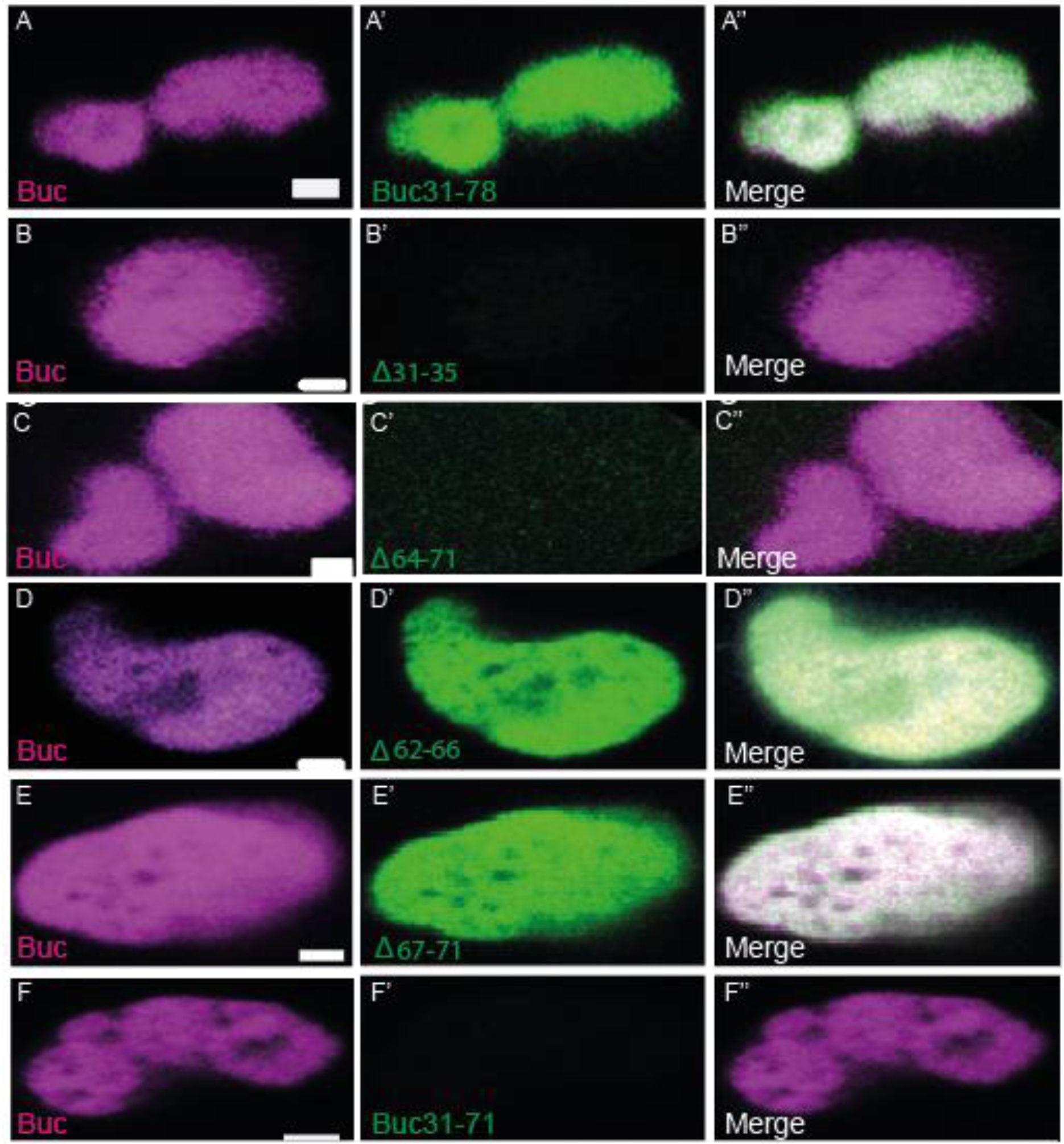
Colocalization of deletion constructs of BucLoc mapping in Figure 5 with Buc-GFP. (A-A’’) shows colocalization of transgenic Buc-GFP (magneta) and BucLoc-m-cherry fusion (aa31-78) (green). (B-B’’) Shows colocalization of transgenic Buc-GFP (magneta) and Buc31-78 (Δ 31-35)-m-cherry fusion (green). (C-C’’) Shows colocalization of transgenic Buc-GFP (magneta) and Buc31-78 (Δ 64-71)-m-cherry fusion (green). (D-D’’) Shows colocalization of transgenic Buc-GFP (magneta) and Buc31-78 (Δ 62-66)-m-cherry fusion (green). (E-E’’) Shows colocalization of transgenic Buc-GFP (magneta) and Buc31-78 (Δ67-71)-m-cherry fusion (green). (F-F’’) Shows colocalization of transgenic Buc-GFP (magneta) and Buc31-71-m-cherry fusion (green). Embryos were injected at 1-cell stage with RNA encoding BucLoc-m-cherry fusions and imaged at high stage. The pictures are representing magnified germ plasm spots. Scale bars: 2 µm.

**Supplementary figure 8.**
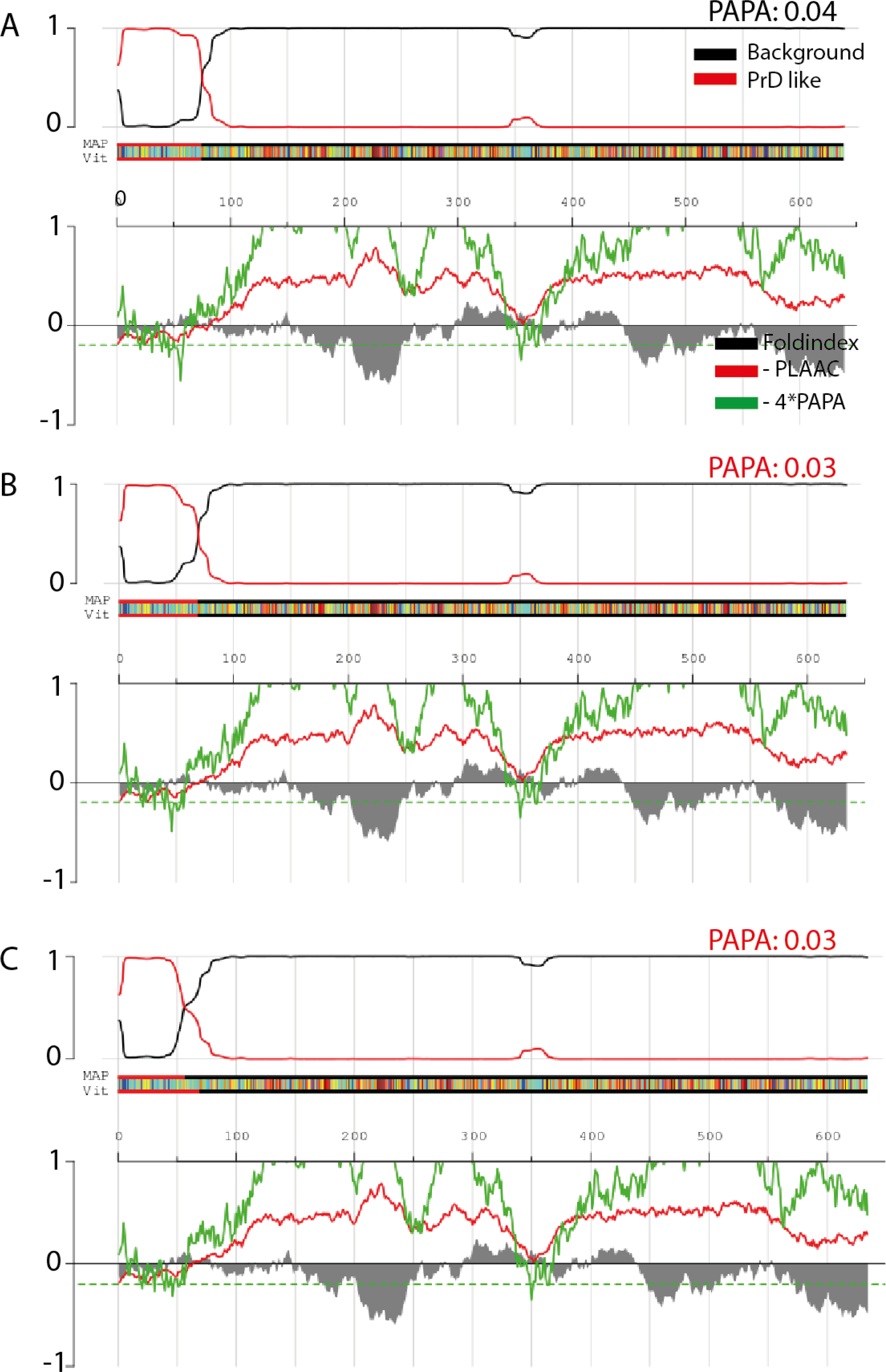
Visualization of PLAAC (Prion-like amino acid composition) outputs. A) WT Buc 74 N-terminal amino acids which are predicted to form aggregates (PAPA 0.04). B) Buc 62-66 (PAPA0.03). C) Buc 67-71 (PAPA 0.03). Upper graph in A, B, and C panels represents a predicted prion like domains (PrDs) (PrD, red line) compared to control (background, black line). Squares with color gradient represent different amino acids in letter code. A second lower graph in A, B, and C panels represents a propensity of a protein to be prion-like protein vs its disorder. FoldIndex (gray) and – PLAAC (red) represent different ways to visualize regions with prion-like composition predicted as disordered. PAPA is a predicted value for amyloid propensity. PAPA multiplied by -4 (-4*PAPA) (green line) is an amyloid prediction value based on a random mutagenesis screen of prion-like proteins. The most negative values of -4*PAPA predicting best the amyloid propensity (threshold is indicated by the green dotted line).

**Supplementary figure 9.**
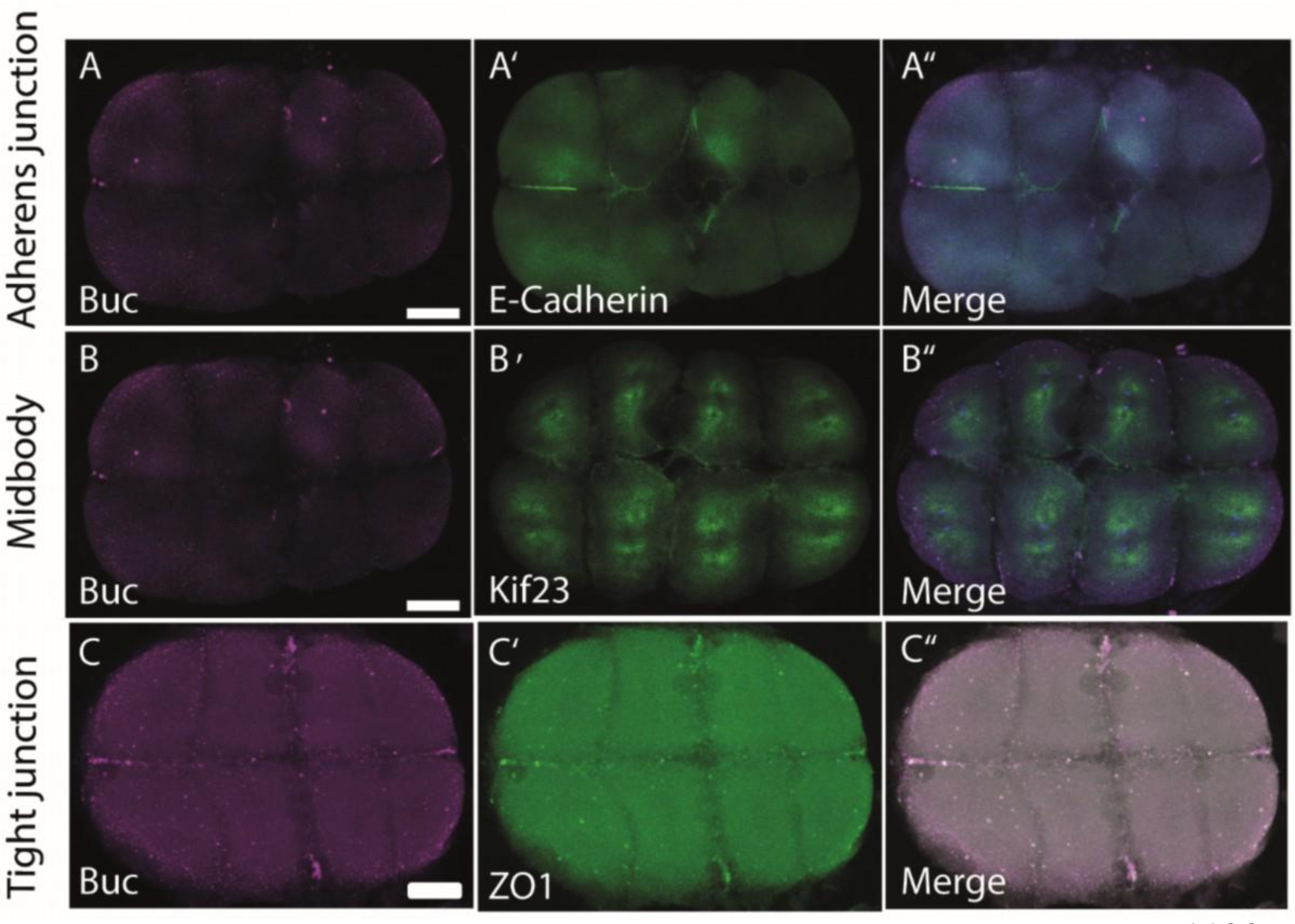
Tight junction protein ZO1 colocalizes with Buc. 1128 Colocalization analysis of Buc with different cellular structure markers. (A, B, C) Show animal view of immunostained 8-cell stage embryos 1^st^ column - Buc (magenta), 2^nd^ column – respective cellular structure (green), 3^rd^ column – merge. (A-A‘‘) Immunostaining for Buc and adherens junction marker E-cadherin; (B-B‘‘) Buc and midbody marker Kif23; (C-C‘‘) Buc and tight junction marker Zonula occludens 1 (ZO1). Scale bars: 50 µm.

**Supplementary figure 10.**
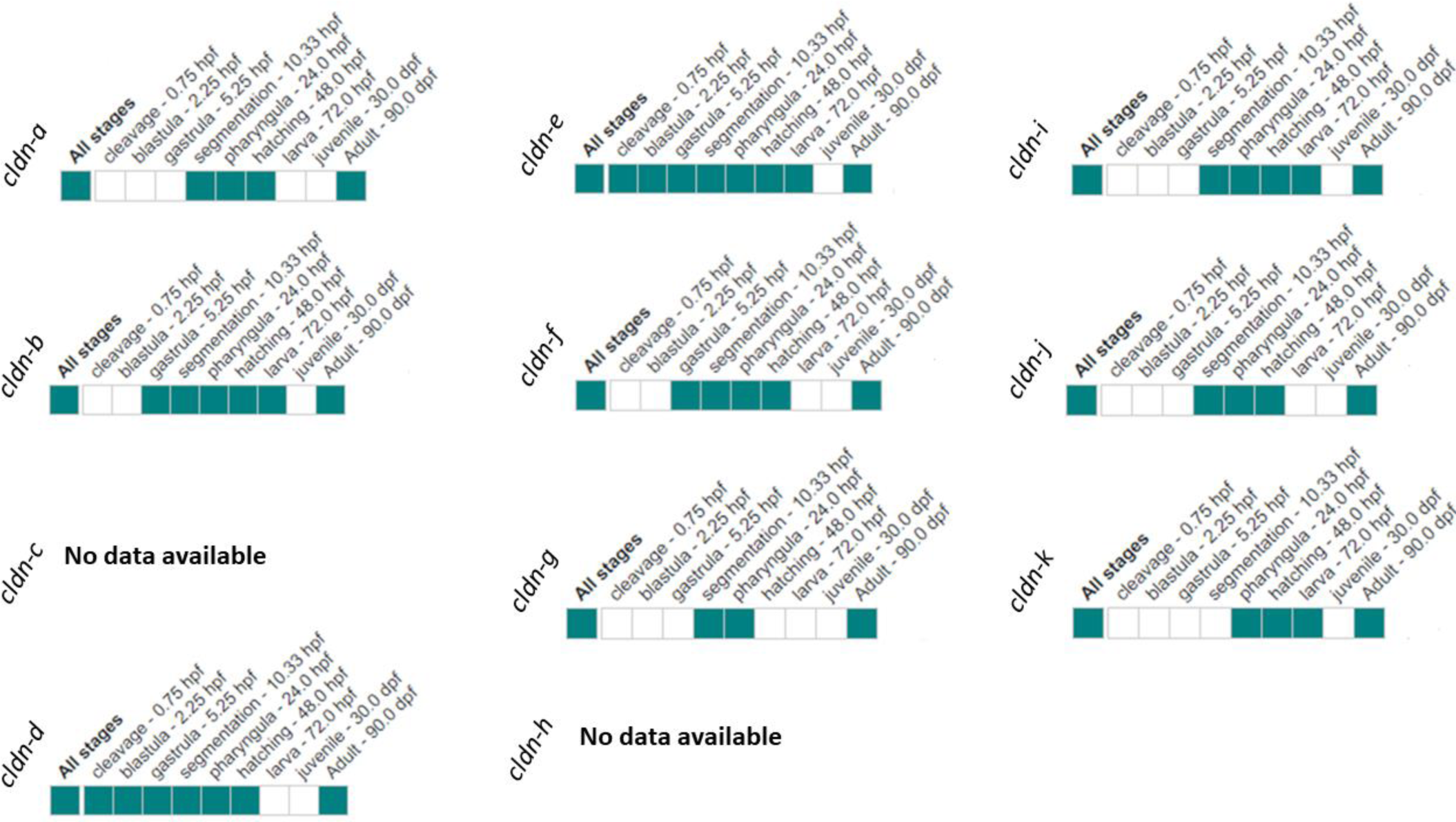
Expression data of zebrafish cldns available on. https://zfin.org/. Expression of cldns at different developmental stages is shown in zebrafish. The green boxes represent annotated expression and the white boxes show no expression. Note only cldn-d and cldn-e showed expression at early developmental stages. The data is updated as on (date February 28th, 2021).

**Supplementary table 1.** . 213 BucLoc interaction candidates selected from 3464 proteins identified by mass spectrometry analysis.

| Identified Protein | Accession Number | GFP Co-IP | Buc-GFP Co-IP | BucLoc-GFP Co-IP |
| --- | --- | --- | --- | --- |
| PREDICTED: protein PRRC2C [Danio rerio] | gi\|528511146 | 79.6 | 1278.6 | 817.9 |
| PREDICTED: zinc finger protein 318-like isoform X1 [Danio rerio] | gi\|528497145 | 12.4 | 666.7 | 512.8 |
| Cluster of PREDICTED: protein PRRC2B isoform X4 [Danio rerio] (gi\|528481247) | gi\|528481247 [2] | 22.6 | 480.1 | 320.0 |
| large proline-rich protein BAT2 [Danio rerio] | gi\|319738640 (+2) | 71.6 | 362.1 | 280.0 |
| PREDICTED: msx2-interacting protein isoform X1 [Danio rerio] | gi\|326678004 (+2) | 63.3 | 434.0 | 257.4 |
| PREDICTED: microtubule-actin cross-linking factor 1, isoforms 1/2/3/5 isoform X1 [Danio rerio] | gi\|528510265 | 0.0 | 546.5 | 218.8 |
| uncharacterized protein LOC792544 [Danio rerio] | gi\|194353937 | 15.8 | 83.1 | 201.9 |
| Cluster of PREDICTED: OTU domain-containing protein 4 isoform X1 [Danio rerio] (gi\|528518829) | gi\|528518829 [2] | 83.8 | 417.6 | 189.9 |
| PREDICTED: YLP motif-containing protein 1 isoform X2 [Danio rerio] | gi\|528511017 | 36.9 | 232.6 | 185.5 |
| PREDICTED: symplekin isoform X2 [Danio rerio] | gi\|326668188 (+1) | 52.9 | 218.7 | 184.5 |
| PREDICTED: microtubule-associated protein futsch-like [Danio rerio] | gi\|528497151 | 1.7 | 123.8 | 181.0 |
| retinoblastoma-binding protein 6 isoform 1 [Danio rerio] | gi\|302632528 (+4) | 13.0 | 75.0 | 159.1 |
| Cluster of PREDICTED: membrane-associated guanylate kinase, WW and PDZ domain-containing protein 1 [Danio rerio] (gi\|528488944) | gi\|528488944 [4] | 62.6 | 191.7 | 157.7 |
| guanine nucleotide-binding protein subunit beta-2-like 1 [Danio rerio] | gi\|18859301 | 53.1 | 208.3 | 149.1 |
| PREDICTED: uncharacterized protein LOC767754 isoform X1 [Danio rerio] | gi\|528467168 (+2) | 45.1 | 104.3 | 134.7 |
| Cluster of eukaryotic translation initiation factor 4A, isoform 1A [Danio rerio] (gi\|38198643) | gi\|38198643 [3] | 43.2 | 230.2 | 127.8 |
| PERQ amino acid-rich with GYF domain-containing protein 2 [Danio rerio] | gi\|71834468 | 55.0 | 261.2 | 123.5 |
| Cluster of zinc finger CCCH domain-containing protein 13 [Danio rerio] (gi\|319738618) | gi\|319738618 | 39.2 | 118.6 | 120.6 |
| Cluster of PREDICTED: cell cycle associated protein 1b isoform X1 [Danio rerio] (gi\|528508316) | gi\|528508316 [3] | 39.1 | 262.3 | 117.6 |
| Cluster of PREDICTED: eukaryotic translation initiation factor 4E transporter isoform X1 [Danio rerio] (gi\|528482894) | gi\|528482894 [2] | 6.5 | 204.5 | 114.7 |
| PREDICTED: cytoskeleton-associated protein 5 isoform X1 [Danio rerio] | gi\|528520895 | 2.9 | 71.0 | 110.6 |
| regulation of nuclear pre-mRNA domain containing 2a [Danio rerio] | gi\|41053979 | 0.0 | 65.7 | 96.8 |
| Cluster of LIM domain only 7b [Danio rerio] (gi\|319996634) | gi\|319996634 [2] | 5.5 | 182.2 | 89.9 |
| GPI-anchored membrane protein 1 [Danio rerio] | gi\|51011059 | 17.0 | 206.9 | 89.8 |
| PREDICTED: trinucleotide repeat-containing gene 6B protein [Danio rerio] | gi\|528495941 | 0.0 | 130.1 | 84.6 |
| Cluster of glutathione S-transferase pi [Danio rerio] (gi\|18858197) | gi\|18858197 | 7.1 | 32.3 | 82.7 |
| Cluster of PREDICTED: pyrroline-5-carboxylate reductase isoform X1 [Danio rerio] (gi\|528495079) | gi\|528495079 [2] | 28.1 | 108.1 | 81.0 |
| ATP synthase subunit O, mitochondrial [Danio rerio] | gi\|51467909 | 4.7 | 21.0 | 76.8 |
| pre-mRNA cleavage complex 2 protein Pcf11 [Danio rerio] | gi\|55925534 | 0.9 | 100.4 | 75.3 |
| PREDICTED: DNA-directed RNA polymerase II subunit RPB1 isoform X2 [Danio rerio] | gi\|528496057 (+1) | 0.0 | 58.5 | 72.8 |
| Cluster of ataxin-2 [Danio rerio] (gi\|190358425) | gi\|190358425 [2] | 30.9 | 149.0 | 70.6 |
| 5'-3' exoribonuclease 1 [Danio rerio] | gi\|289577074 (+1) | 1.4 | 114.6 | 65.5 |
| voltage-dependent anion-selective channel protein 2 [Danio rerio] | gi\|41054601 (+1) | 16.5 | 33.7 | 65.5 |
| Cluster of splicing factor 45 [Danio rerio] (gi\|41055474) | gi\|41055474 [2] | 13.5 | 132.2 | 57.6 |
| voltage-dependent anion-selective channel protein 1 [Danio rerio] | gi\|47777306 | 17.5 | 42.1 | 57.0 |
| PREDICTED: protein FAM208A [Danio rerio] | gi\|528517763 | 1.9 | 70.5 | 54.5 |
| single-stranded DNA-binding protein, mitochondrial [Danio rerio] | gi\|62955585 | 11.5 | 24.0 | 54.4 |
| PREDICTED: tyrosine 3-monooxygenase/tryptophan 5-monooxygenase activation protein, zeta polypeptide isoform X1 [Danio rerio] | gi\|528509453 | 3.6 | 13.6 | 53.7 |
| non-POU domain-containing octamer-binding protein [Danio rerio] | gi\|42415509 | 23.4 | 96.3 | 52.9 |
| Cluster of PREDICTED: fish-egg lectin isoform X1 [Danio rerio] (gi\|528491480) | gi\|528491480 [5] | 7.2 | 64.2 | 52.2 |
| alpha-2-macroglobulin-like precursor [Danio rerio] | gi\|320118891 | 0.0 | 17.3 | 50.3 |
| Cluster of PREDICTED: chromodomain helicase DNA binding protein 4 isoform X1 [Danio rerio] (gi\|528509046) | gi\|528509046 [4] | 1.7 | 30.1 | 49.5 |
| 14-3-3 protein epsilon [Danio rerio] | gi\|47086819 | 4.6 | 12.4 | 47.8 |
| Cluster of uncharacterized protein LOC100141336 precursor [Danio rerio] (gi\|168823478) | gi\|168823478 [5] | 11.2 | 41.2 | 47.5 |
| PREDICTED: protein SCAF11 isoform X1 [Danio rerio] | gi\|528475676 (+2) | 0.9 | 52.1 | 45.5 |
| PREDICTED: cytoplasmic dynein 2 heavy chain 1-like, partial [Danio rerio] | gi\|528502710 | 20.8 | 42.4 | 44.2 |
| Cluster of PREDICTED: pericentriolar material 1 protein isoform X7 [Danio rerio] (gi\|528467744) | gi\|528467744 [3] | 1.7 | 12.9 | 44.1 |
| alpha-2-macroglobulin-like [Danio rerio] | gi\|319655740 (+2) | 2.9 | 18.7 | 43.1 |
| PREDICTED: ubiquitin carboxyl-terminal hydrolase 10 isoformX1 [Danio rerio] | gi\|326669691 | 1.0 | 138.2 | 42.9 |
| PREDICTED: cleavage stimulation factor subunit 2-like isoform X1 [Danio rerio] | gi\|326673799 | 11.3 | 86.7 | 40.3 |
| Cluster of PREDICTED: alpha-2-macroglobulin isoformX1 [Danio rerio] (gi\|326665588) | gi\|326665588 [2] | 8.9 | 60.0 | 39.8 |
| PREDICTED: eukaryotic translation initiation factor 4 gamma 3 [Danio rerio] | gi\|528517986 | 3.0 | 151.5 | 39.8 |
| Ndufa9 protein [Danio rerio] | gi\|157423514 (+2) | 9.6 | 22.4 | 39.6 |
| Zgc:158157 protein [Danio rerio] | gi\|55250357 | 0.0 | 63.4 | 39.5 |
| Cluster of myosin light chain alkali, smooth-muscle isoform [Danio rerio] (gi\|47174755) | gi\|47174755 [6] | 8.7 | 17.3 | 39.1 |
| Cluster of PREDICTED: uncharacterized protein LOC100000125 isoform X2 [Danio rerio] (gi\|528510415) | gi\|528510415 [3] | 6.1 | 103.3 | 38.8 |
| signal-induced proliferation-associated 1-like protein 1 [Danio rerio] | gi\|529250122 | 2.8 | 31.4 | 38.7 |
| eukaryotic translation initiation factor 4E-1B [Danio rerio] | gi\|18858611 | 11.8 | 92.6 | 37.4 |
| Cyc1 protein [Danio rerio] | gi\|51327354 (+1) | 11.5 | 32.9 | 35.8 |
| complement component 1 Q subcomponent-binding protein, mitochondrial [Danio rerio] | gi\|324021711 (+1) | 9.0 | 19.2 | 35.8 |
| LOC449616 protein [Danio rerio] | gi\|213624848 (+2) | 0.0 | 92.5 | 35.8 |
| PREDICTED: cyclin-dependent kinase 13 [Danio rerio] | gi\|326679472 | 5.8 | 67.7 | 33.7 |
| Cluster of CWF19-like protein 2 [Danio rerio] (gi\|76253886) | gi\|76253886 [2] | 0.0 | 28.5 | 32.6 |
| nanog homeobox [Danio rerio] | gi\|528505177 (+1) | 3.9 | 60.8 | 32.4 |
| PREDICTED: zinc finger CCCH domain-containing protein 4-like isoform X1 [Danio rerio] | gi\|528501822 | 0.0 | 18.3 | 31.6 |
| Cluster of PREDICTED: unconventional myosin-Va [Danio rerio] (gi\|326680074) | gi\|326680074 [4] | 3.7 | 20.9 | 31.4 |
| PREDICTED: protein FAM208A-like [Danio rerio] | gi\|528516938 | 0.0 | 23.0 | 31.2 |
| glutamate dehydrogenase 1b [Danio rerio] | gi\|41282194 | 0.0 | 27.4 | 31.0 |
| S-phase kinase-associated protein 1 [Danio rerio] | gi\|41152201 | 14.5 | 29.8 | 30.8 |
| PREDICTED: tyrosine-protein phosphatase non-receptor type 13 isoform X1 [Danio rerio] | gi\|528513092 (+4) | 6.4 | 107.4 | 30.7 |
| Cluster of ADP-ribosylation factor 1 like [Danio rerio] (gi\|41393117) | gi\|41393117 [2] | 0.0 | 11.2 | 30.1 |
| Si:dkey-16k6.1 protein [Danio rerio] | gi\|45767805 (+1) | 2.4 | 7.8 | 29.8 |
| Cluster of ras homolog gene family, member Ad [Danio rerio] (gi\|50539958) | gi\|50539958 [3] | 1.2 | 16.1 | 29.4 |
| tight junction protein ZO-2 isoform 1 [Danio rerio] | gi\|320118869 (+2) | 0.0 | 13.5 | 29.2 |
| Cluster of uncharacterized protein LOC569235 precursor [Danio rerio] (gi\|350536793) | gi\|350536793 [3] | 13.0 | 42.7 | 29.2 |
| ATP-dependent RNA helicase DDX42 [Danio rerio] | gi\|302318882 | 0.0 | 73.3 | 27.9 |
| Cluster of PREDICTED: hypothetical protein LOC565404 [Danio rerio] (gi\|189517232) | gi\|189517232 [5] | 6.7 | 58.2 | 27.5 |
| PREDICTED: telomerase-binding protein EST1A-like isoform X1 [Danio rerio] | gi\|326671361 (+1) | 0.0 | 90.3 | 27.4 |
| PREDICTED: ATP synthase subunit b, mitochondrial isoform X1 [Danio rerio] | gi\|528487650 (+1) | 3.5 | 9.1 | 27.1 |
| mitochondrial import inner membrane translocase subunit Tim13 [Danio rerio] | gi\|50539998 | 0.0 | 10.2 | 26.9 |
| PREDICTED: RNA-binding protein 27 isoform X2 [Danio rerio] | gi\|528491090 (+1) | 5.8 | 32.9 | 26.7 |
| PREDICTED: nudC domain-containing protein 1 isoform X1 [Danio rerio] | gi\|528509803 | 2.7 | 11.2 | 26.6 |
| regulation of nuclear pre-mRNA domain-containing protein 1B [Danio rerio] | gi\|41054665 (+3) | 0.0 | 63.3 | 25.4 |
| nuclear pore complex protein Nup133 [Danio rerio] | gi\|47087231 | 0.0 | 8.9 | 25.1 |
| PREDICTED: RNA-binding protein 6 isoform X1 [Danio rerio] | gi\|528483145 | 0.0 | 25.4 | 24.6 |
| Cluster of PREDICTED: RNA-binding protein 26 [Danio rerio] (gi\|528481486) | gi\|528481486 [2] | 2.5 | 30.2 | 24.6 |
| glutamate dehydrogenase 1a [Danio rerio] | gi\|47086875 | 0.0 | 11.1 | 24.5 |
| Cluster of Bub3 protein [Danio rerio] (gi\|53734038) | gi\|53734038 [2] | 3.2 | 16.2 | 23.8 |
| Cluster of PREDICTED: cat eye syndrome critical region protein 2 isoform X2 [Danio rerio] (gi\|528521383) | gi\|528521383 [2] | 0.0 | 46.3 | 23.6 |
| LYR motif-containing protein 4 [Danio rerio] | gi\|256000753 | 0.0 | 7.7 | 23.4 |
| mitochondrial inner membrane protein [Danio rerio] | gi\|47777298 (+1) | 3.5 | 15.4 | 22.5 |
| Abcf2 protein [Danio rerio] | gi\|42542861 (+1) | 7.3 | 27.2 | 22.5 |
| DNA-directed RNA polymerase II subunit RPB2 [Danio rerio] | gi\|302488402 | 0.0 | 33.2 | 22.5 |
| Cluster of SNW domain-containing protein 1 [Danio rerio] (gi\|50838798) | gi\|50838798 | 7.9 | 39.3 | 22.4 |
| PREDICTED: NFX1-type zinc finger-containing protein 1-like [Danio rerio] | gi\|528483584 | 5.8 | 39.3 | 22.4 |
| periphilin-1 [Danio rerio] | gi\|121583944 (+1) | 1.4 | 33.7 | 22.3 |
| PREDICTED: unconventional myosin-IXa isoform X1 [Danio rerio] | gi\|528486090 (+1) | 0.0 | 56.0 | 22.2 |
| PREDICTED: histone-lysine N-methyltransferase SETD1A isoform X1 [Danio rerio] | gi\|326666050 | 0.0 | 38.1 | 21.9 |
| PREDICTED: death-inducer obliterator 1-like [Danio rerio] | gi\|528516988 | 0.0 | 59.7 | 21.9 |
| Cluster of LOC100000597 protein [Danio rerio] (gi\|66910514) | gi\|66910514 [5] | 0.0 | 10.2 | 21.7 |
| dolichyl-diphosphooligosaccharide--protein glycosyltransferase subunit DAD1 [Danio rerio] | gi\|154426290 | 0.0 | 10.2 | 21.1 |
| protein QIL1 [Danio rerio] | gi\|113678245 | 4.2 | 10.2 | 21.1 |
| PREDICTED: E3 ubiquitin-protein ligase TTC3 isoform X2 [Danio rerio] | gi\|528491184 | 2.8 | 70.6 | 21.0 |
| serine/threonine-protein phosphatase 2A 56 kDa regulatory subunit delta isoform [Danio rerio] | gi\|50726892 | 0.0 | 5.5 | 20.9 |
| cytochrome b-c1 complex subunit Rieske, mitochondrial [Danio rerio] | gi\|157073897 | 0.0 | 6.1 | 20.7 |
| Cluster of Protein regulator of cytokinesis 1 [Danio rerio] (gi\|28279644) | gi\|28279644 [2] | 7.5 | 61.7 | 20.4 |
| PREDICTED: E3 ubiquitin-protein ligase HERC2, partial [Danio rerio] | gi\|528483182 | 0.0 | 8.5 | 20.0 |
| Cluster of LOC563225 protein [Danio rerio] (gi\|115292012) | gi\|115292012 [2] | 7.9 | 17.8 | 19.7 |
| Cluster of eukaryotic translation initiation factor 4B [Danio rerio] (gi\|47550837) | gi\|47550837 [6] | 0.0 | 41.9 | 19.6 |
| high mobility group protein B2 [Danio rerio] | gi\|82658290 | 0.9 | 5.7 | 19.4 |
| Icln protein [Danio rerio] | gi\|44890532 (+2) | 7.6 | 23.6 | 19.3 |
| nuclear pore complex protein Nup160 [Danio rerio] | gi\|41054908 | 0.0 | 5.3 | 19.3 |
| Cluster of TNF receptor-associated protein 1 [Danio rerio] (gi\|165972373) | gi\|165972373 [2] | 0.0 | 71.3 | 18.8 |
| Cluster of PREDICTED: microtubule-associated serine/threonine-protein kinase 3-like [Danio rerio] (gi\|528470977) | gi\|528470977 [2] | 0.0 | 25.5 | 18.8 |
| M-phase phosphoprotein 8 [Danio rerio] | gi\|187607764 | 2.4 | 42.2 | 18.7 |
| PREDICTED: PHD finger protein 3 isoform X1 [Danio rerio] | gi\|528498598 (+1) | 0.0 | 18.4 | 18.5 |
| PREDICTED: sideroflexin-3 isoform X1 [Danio rerio] | gi\|528497698 | 1.9 | 7.2 | 18.2 |
| Cluster of Epithelial cell adhesion molecule [Danio rerio] (gi\|44890710) | gi\|44890710 [2] | 1.8 | 13.9 | 18.0 |
| Sept2 protein [Danio rerio] | gi\|115313325 (+3) | 0.0 | 13.3 | 17.6 |
| Cluster of PREDICTED: regulation of nuclear pre-mRNA domain-containing protein 2 isoform X1 [Danio rerio] (gi\|528502856) | gi\|528502856 [2] | 0.0 | 12.6 | 17.5 |
| Zgc:77560 protein [Danio rerio] | gi\|42542976 (+1) | 2.4 | 66.6 | 17.4 |
| Cluster of PREDICTED: nucleolar pre-ribosomal-associated protein 1-like [Danio rerio] (gi\|528501168) | gi\|528501168 [2] | 2.9 | 20.9 | 17.3 |
| ras-related protein Rab-14 [Danio rerio] | gi\|41393147 | 0.0 | 5.1 | 16.8 |
| NADH dehydrogenase [ubiquinone] 1 beta subcomplex subunit 10 [Danio rerio] | gi\|41152268 | 3.5 | 17.2 | 16.8 |
| Cluster of oxoglutarate (alpha-ketoglutarate) dehydrogenase (lipoamide) [Danio rerio] (gi\|254028264) | gi\|254028264 [2] | 0.0 | 5.6 | 16.6 |
| Pard3 protein [Danio rerio] | gi\|190339230 (+4) | 0.9 | 32.3 | 16.6 |
| DNA-directed RNA polymerase II subunit RPB3 [Danio rerio] | gi\|269784633 | 0.0 | 22.8 | 16.3 |
| RecName: Full=Protein CASC3; AltName: Full=Cancer susceptibility candidate gene 3 protein homolog; AltName: Full=Metastatic lymph node protein 51 homolog; Short=DrMLN51; Short=Protein MLN 51 homolog | gi\|123886565 (+1) | 6.3 | 28.9 | 16.1 |
| Cluster of serine/threonine kinase 36 (fused homolog, Drosophila) [Danio rerio] (gi\|320043268) | gi\|320043268 | 6.0 | 13.9 | 16.0 |
| cytotoxic granule-associated RNA binding protein 1 [Danio rerio] | gi\|47086779 | 6.4 | 16.9 | 16.0 |
| Cluster of PREDICTED: uncharacterized protein LOC393431 isoform X1 [Danio rerio] (gi\|528486792) | gi\|528486792 | 0.0 | 9.1 | 16.0 |
| Cluster of nuclear receptor corepressor 2 [Danio rerio] (gi\|380420327) | gi\|380420327 [4] | 0.0 | 57.7 | 15.9 |
| Cas-Br-M (murine) ecotropic retroviral transforming sequence-like 1 [Danio rerio] | gi\|41055074 | 3.3 | 20.8 | 15.7 |
| PREDICTED: polyribonucleotide 5'-hydroxyl-kinase Clp1-like [Danio rerio] | gi\|189528302 | 0.0 | 53.2 | 15.4 |
| uncharacterized protein LOC556124 [Danio rerio] | gi\|157909776 | 1.4 | 30.5 | 15.2 |
| succinate dehydrogenase [ubiquinone] iron-sulfur subunit, mitochondrial precursor [Danio rerio] | gi\|148922926 | 5.4 | 11.3 | 14.9 |
| Cluster of cAMP-dependent protein kinase catalytic subunit alpha [Danio rerio] (gi\|130493522) | gi\|130493522 [3] | 0.0 | 8.8 | 14.9 |
| serine/threonine-protein kinase 3 [Danio rerio] | gi\|41054445 (+1) | 3.6 | 49.0 | 14.8 |
| uncharacterized protein LOC541537 [Danio rerio] | gi\|62122901 | 0.0 | 7.4 | 14.6 |
| chromodomain-helicase-DNA-binding protein 8 [Danio rerio] | gi\|320461545 (+2) | 0.0 | 22.7 | 14.6 |
| PREDICTED: splicing factor, arginine/serine-rich 15 [Danio rerio] | gi\|528492333 | 0.0 | 20.4 | 14.4 |
| zinc finger HIT domain-containing protein 3 [Danio rerio] | gi\|41053670 | 3.5 | 33.1 | 14.0 |
| eukaryotic translation initiation factor 6 [Danio rerio] | gi\|41055624 (+2) | 0.0 | 9.1 | 13.9 |
| PREDICTED: nuclear pore complex protein Nup153 isoform X1 [Danio rerio] | gi\|528510621 | 0.0 | 17.5 | 13.7 |
| PREDICTED: uncharacterized protein LOC436879 isoform X4 [Danio rerio] | gi\|528517594 | 0.0 | 11.1 | 13.3 |
| Zgc:111960 protein [Danio rerio] | gi\|166796880 | 0.0 | 11.6 | 13.2 |
| very low-density lipoprotein receptor precursor [Danio rerio] | gi\|169646705 (+4) | 0.0 | 5.1 | 13.0 |
| C-terminal binding protein 1 [Danio rerio] | gi\|40254690 | 3.3 | 11.6 | 12.9 |
| claudin-like protein ZF-A89 [Danio rerio] | gi\|30725822 | 0.0 | 5.2 | 12.8 |
| Cluster of PREDICTED: C2 domain-containing protein 3 [Danio rerio] (gi\|528501432) | gi\|528501432 | 0.0 | 9.2 | 12.7 |
| PREDICTED: histone deacetylase 1 isoform X1 [Danio rerio] | gi\|528510099 | 2.6 | 29.6 | 12.0 |
| peptidyl-prolyl cis-trans isomerase-like 1 [Danio rerio] | gi\|77683061 | 1.9 | 18.6 | 11.9 |
| Cluster of ras-related C3 botulinum toxin substrate 1 [Danio rerio] (gi\|54792776) | gi\|54792776 [2] | 3.5 | 11.7 | 11.9 |
| PREDICTED: uncharacterized protein KIAA0556-like [Danio rerio] | gi\|528520519 | 0.0 | 14.9 | 11.8 |
| PREDICTED: trinucleotide repeat-containing gene 6B protein-like isoform X2 [Danio rerio] | gi\|528472879 (+1) | 0.0 | 23.4 | 11.7 |
| nuclear pore complex protein Nup107 [Danio rerio] | gi\|71834480 | 1.8 | 6.9 | 11.3 |
| Cluster of PREDICTED: tight junction protein ZO-1-like [Danio rerio] (gi\|528521995) | gi\|528521995 | 0.0 | 12.5 | 11.3 |
| DIS3-like exonuclease 1 [Danio rerio] | gi\|160333118 | 0.0 | 32.6 | 11.3 |
| PREDICTED: PAB-dependent poly(A)-specific ribonuclease subunit 2-like isoform X2 [Danio rerio] | gi\|528518391 (+1) | 0.0 | 13.3 | 11.2 |
| Cluster of PREDICTED: PERQ amino acid-rich with GYF domain-containing protein 1-like isoform X1 [Danio rerio] (gi\|528492127) | gi\|528492127 [2] | 0.0 | 13.3 | 11.2 |
| uncharacterized protein LOC550263 [Danio rerio] | gi\|62955177 | 0.0 | 14.9 | 11.2 |
| Cluster of PREDICTED: serine/threonine-protein kinase TAO2-like [Danio rerio] (gi\|125812164) | gi\|125812164 | 0.0 | 17.9 | 10.7 |
| PREDICTED: uncharacterized protein LOC503771 isoform X3 [Danio rerio] | gi\|528519733 | 0.0 | 17.1 | 10.6 |
| Cluster of LOC559853 protein, partial [Danio rerio] (gi\|79151969) | gi\|79151969 [2] | 0.0 | 38.9 | 10.4 |
| PREDICTED: serine/threonine-protein kinase LATS1 isoform X1 [Danio rerio] | gi\|528510786 | 0.0 | 20.4 | 10.2 |
| PREDICTED: cyclin-dependent kinase 12 [Danio rerio] | gi\|528509066 | 0.9 | 29.5 | 10.0 |
| poly(rC)-binding protein 2 [Danio rerio] | gi\|41055221 | 0.0 | 12.7 | 9.9 |
| nanog homeobox [Danio rerio] | gi\|148357118 | 0.0 | 28.5 | 9.9 |
| Cluster of TIA1 cytotoxic granule-associated RNA binding protein [Danio rerio] (gi\|37681959) | gi\|37681959 [4] | 4.8 | 33.1 | 9.9 |
| protein mago nashi homolog [Danio rerio] | gi\|62955377 | 3.8 | 15.8 | 9.8 |
| nucleoporin 98 [Danio rerio] | gi\|320118905 (+1) | 0.0 | 7.5 | 9.7 |
| uncharacterized protein LOC100126100 precursor [Danio rerio] | gi\|157954446 (+1) | 2.9 | 22.5 | 9.7 |
| N-acetyltransferase 10 [Danio rerio] | gi\|41055301 | 4.7 | 15.0 | 9.7 |
| OCIA domain-containing protein 1 [Danio rerio] | gi\|41053513 (+1) | 0.0 | 5.8 | 9.5 |
| cytochrome c oxidase subunit II [Danio rerio] | gi\|8395615 | 2.4 | 5.1 | 9.5 |
| PREDICTED: transcription factor 19 [Danio rerio] | gi\|292624089 | 0.0 | 17.6 | 9.4 |
| signal recognition particle 9 [Danio rerio] | gi\|41055367 | 0.0 | 5.1 | 9.4 |
| actin related protein 2/3 complex subunit 4 [Danio rerio] | gi\|45387521 | 0.0 | 7.4 | 9.1 |
| PREDICTED: lysine-specific demethylase 6A isoform X1 [Danio rerio] | gi\|528490350 (+1) | 0.0 | 26.6 | 8.8 |
| exosome complex exonuclease RRP4 [Danio rerio] | gi\|339717151 | 2.9 | 22.8 | 8.8 |
| PREDICTED: bromodomain adjacent to zinc finger domain, 2A isoform X1 [Danio rerio] | gi\|528517226 (+1) | 0.0 | 8.0 | 8.7 |
| PREDICTED: AF4/FMR2 family member 4 isoform X1 [Danio rerio] | gi\|528514500 (+3) | 2.6 | 13.3 | 8.6 |
| Cluster of PREDICTED: kinesin family member 13A [Danio rerio] (gi\|528505240) | gi\|528505240 [5] | 0.0 | 5.4 | 8.4 |
| exosome complex exonuclease RRP45 [Danio rerio] | gi\|54400656 | 0.0 | 20.2 | 8.4 |
| exosome complex component MTR3 [Danio rerio] | gi\|66472734 | 1.0 | 14.4 | 8.1 |
| MGC174638 protein [Danio rerio] | gi\|156230391 (+2) | 0.0 | 7.8 | 8.0 |
| Cluster of cyclin-L1 [Danio rerio] (gi\|41054323) | gi\|41054323 | 3.6 | 13.3 | 7.7 |
| Xrcc5 protein [Danio rerio] | gi\|133777834 | 0.0 | 12.9 | 7.7 |
| sorting and assembly machinery component 50 homolog B [Danio rerio] | gi\|55925219 | 0.0 | 5.2 | 7.4 |
| Cluster of PREDICTED: rho GTPase-activating protein 21 isoform X1 [Danio rerio] (gi\|528470502) | gi\|528470502 [2] | 0.0 | 10.9 | 7.4 |
| 60S ribosomal protein L29 [Danio rerio] | gi\|51010951 | 2.9 | 11.6 | 7.1 |
| immediate early response 3-interacting protein 1 precursor [Danio rerio] | gi\|356991159 | 0.0 | 5.1 | 7.0 |
| transmembrane and coiled-coil domains 1 [Danio rerio] | gi\|50540216 (+1) | 1.9 | 8.7 | 7.0 |
| PREDICTED: wu:fc48e01 [Danio rerio] | gi\|125820176 | 0.0 | 18.4 | 6.8 |
| Zgc:123096 protein, partial [Danio rerio] | gi\|50417024 (+1) | 0.0 | 6.1 | 6.6 |
| PREDICTED: protein TANC1-like isoform X2 [Danio rerio] | gi\|528481904 (+1) | 0.0 | 9.1 | 6.4 |
| Centrin2 [Danio rerio] | gi\|161213715 (+1) | 1.9 | 14.9 | 6.3 |
| PREDICTED: dehydrogenase/reductase SDR family member 7B isoform X1 [Danio rerio] | gi\|528473478 | 0.0 | 10.8 | 6.3 |
| cyclin T2b [Danio rerio] | gi\|47086855 | 2.7 | 16.8 | 6.3 |
| U4/U6.U5 tri-snRNP-associated protein 1 [Danio rerio] | gi\|50540414 | 0.0 | 15.6 | 5.9 |
| PREDICTED: uveal autoantigen with coiled-coil domains and ankyrin repeats [Danio rerio] | gi\|292616084 (+1) | 0.0 | 6.6 | 5.9 |
| C-Myc-binding protein [Danio rerio] | gi\|91176306 | 0.0 | 15.4 | 5.9 |
| transcription elongation factor B polypeptide 1 [Danio rerio] | gi\|52219182 (+1) | 0.0 | 7.7 | 5.9 |
| Cluster of cytochrome c oxidase assembly factor 5 [Danio rerio] (gi\|238859533) | gi\|238859533 | 0.0 | 5.1 | 5.9 |
| PREDICTED: uncharacterized protein DKFZp762I1415-like [Danio rerio] | gi\|68399071 | 0.0 | 5.1 | 5.9 |
| Ku70 autoantigen [Danio rerio] | gi\|114215700 (+2) | 1.6 | 7.4 | 5.7 |
| uncharacterized protein LOC100216070 [Xenopus (Silurana) tropicalis] | gi\|213983243 (+1) | 0.0 | 11.3 | 5.6 |
| RNA-binding protein 8A [Danio rerio] | gi\|61651846 | 2.4 | 9.0 | 5.6 |
| G kinase anchoring protein 1 [Danio rerio] | gi\|32451811 (+3) | 0.0 | 17.8 | 5.6 |
| PREDICTED: nuclear receptor coactivator 3 [Danio rerio] | gi\|326671802 | 0.0 | 14.8 | 5.3 |
| Cluster of PREDICTED: mps one binder kinase activator-like 1B-like [Danio rerio] (gi\|292614396) | gi\|292614396 [5] | 0.0 | 9.1 | 5.1 |
| coiled-coil-helix-coiled-coil-helix domain-containing protein 2, mitochondrial [Danio rerio] | gi\|41152140 | 0.0 | 8.1 | 5.1 |
| store-operated calcium entry-associated regulatory factor precursor [Danio rerio] | gi\|115495791 (+2) | 0.0 | 13.2 | 5.0 |

**Supplementary table 2.**
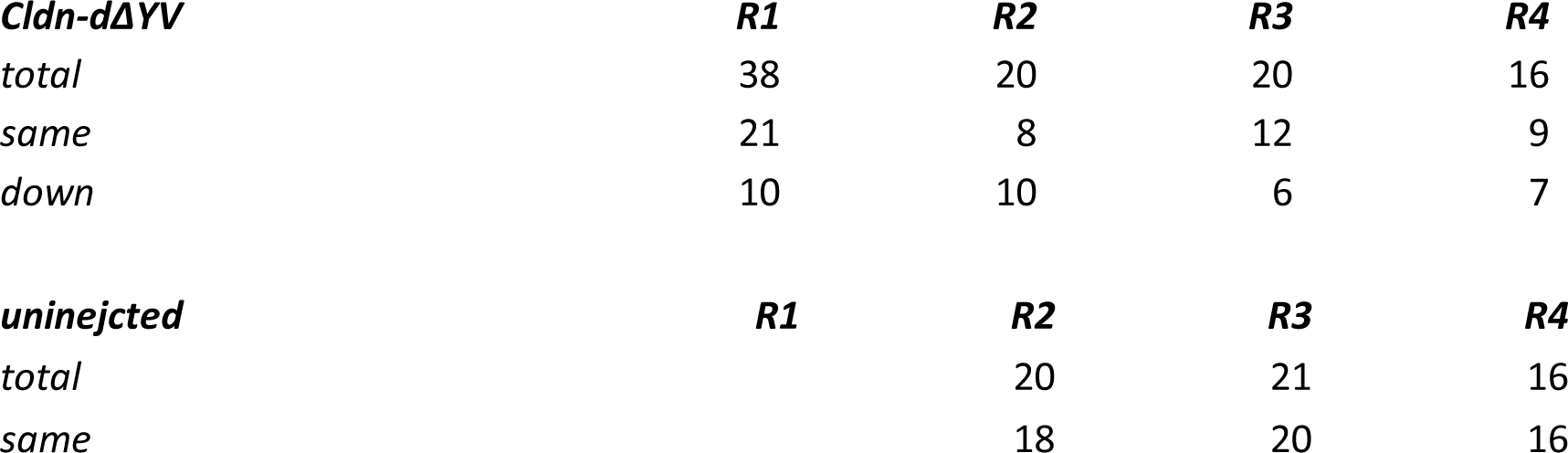
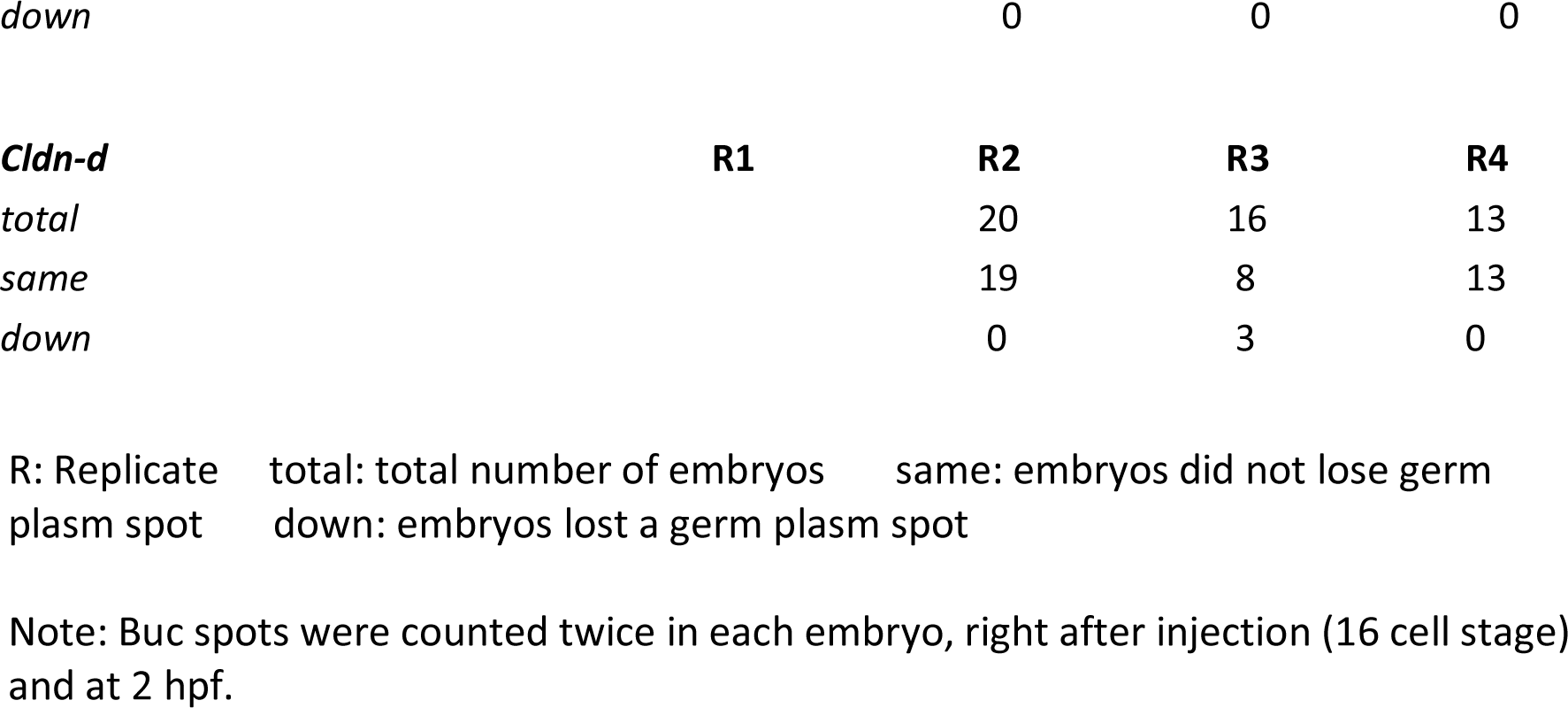
. Data for 16-cell injection assay.

Digital appendix

For a full list of the 3464 proteins identified by mass spectrometry analysis please open the Excel file.

